# Dilated cardiomyopathy mutation in beta-cardiac myosin enhances actin activation of the power stroke and phosphate release

**DOI:** 10.1101/2023.11.10.566646

**Authors:** Skylar M. L. Bodt, Jinghua Ge, Wen Ma, David V. Rasicci, Rohini Desetty, J. Andrew McCammon, Christopher M. Yengo

**Affiliations:** Department of Cellular and Molecular Physiology, Pennsylvania State University College of Medicine, Hershey, Pennsylvania; Department of Chemistry and Biochemistry, University of California San Diego, San Diego, California

**Keywords:** Myosin, Actin, Heart Failure, Muscle Contraction, Enzymology

## Abstract

Inherited mutations in human beta-cardiac myosin (M2β) can lead to severe forms of heart failure. The E525K mutation in M2β is associated with dilated cardiomyopathy (DCM) and was found to stabilize the interacting heads motif (IHM) and autoinhibited super-relaxed (SRX) state in dimeric heavy meromyosin. However, in monomeric M2β subfragment 1 (S1) we found that E525K enhances (3-fold) the maximum steady-state actin-activated ATPase activity (*k*_cat_) and decreases (6-fold) the actin concentration at which ATPase is one-half maximal (*K*_ATPase_). We also found a 3 to 4-fold increase in the actin-activated power stroke and phosphate release rate constants at 30 μM actin, which overall enhanced the duty ratio 3-fold. Loaded motility assays revealed that the enhanced intrinsic motor activity translates to increased ensemble force in M2β S1. Glutamate 525, located near the actin binding region in the so-called activation loop, is highly conserved and predicted to form a salt-bridge with another conserved residue (lysine 484) in the relay helix. Enhanced sampling molecular dynamics simulations predict that the charge reversal mutation disrupts the E525-K484 salt-bridge, inducing conformations with a more flexible relay helix and a wide phosphate release tunnel. Our results highlight a highly conserved allosteric pathway associated with actin activation of the power stroke and phosphate release and suggest an important feature of the autoinhibited IHM is to prevent this region of myosin from interacting with actin. The ability of the E525K mutation to stabilize the IHM likely overrides the enhanced intrinsic motor properties, which may be key to triggering DCM pathogenesis.

**Significance Statement:** Heart disease can be caused by inherited mutations in beta-cardiac myosin, the molecular motor that powers systolic contraction in the ventricles of the heart. However, it remains unclear how these mutations lead to contractile dysfunction and pathogenic remodeling of the heart. We investigated a unique dilated cardiomyopathy mutation (E525K) that dramatically stabilizes the autoinhibited state while enhancing intrinsic motor function. Thus, we examined how this mutation impacts transient kinetic steps of the ATPase cycle, motile properties, and structural changes associated with the power stroke and phosphate release. Our results provide a kinetic and structural basis for how beta-cardiac myosin mutations may disrupt molecular-level contractile function in complex ways, which may inform the development of targeted therapeutics.

## Introduction

Human beta-cardiac myosin (M2β), is the molecular motor that powers systolic contraction within the heart by utilizing a conserved actomyosin ATPase mechanism (**Fig. 1**) (1, 2). It is well established that generation of force and power in cardiac muscle is determined by the recruitment of myosin motors during the systolic rise of calcium. During cardiac systole, cytoplasmic calcium binds troponin-C causing a conformational shift in tropomyosin that allows myosin to bind actin (3). Myosin heads drive contraction according to the sliding filament theory by harnessing mechanical energy from ATP hydrolysis to initiate the force generating power stroke (4). More recently it has been revealed that muscle myosins can occupy an autoinhibited state called the super relaxed (SRX) state with 10-fold slower ATP turnover that plays an important role in thick filament regulation (5-9). Reports have found correlations between this SRX state and the interacting heads motif (IHM), a conformation where myosin heads fold back on a region of the tail (subfragment 2) and interact with each other (10). The intricate balance between the force generating mechanism and autoinhibition of the thick filament remains elusive (11). Examining these biochemical and structural events is crucial for understanding the system of energy transduction and regulation in myosin.

**Figure 1.**
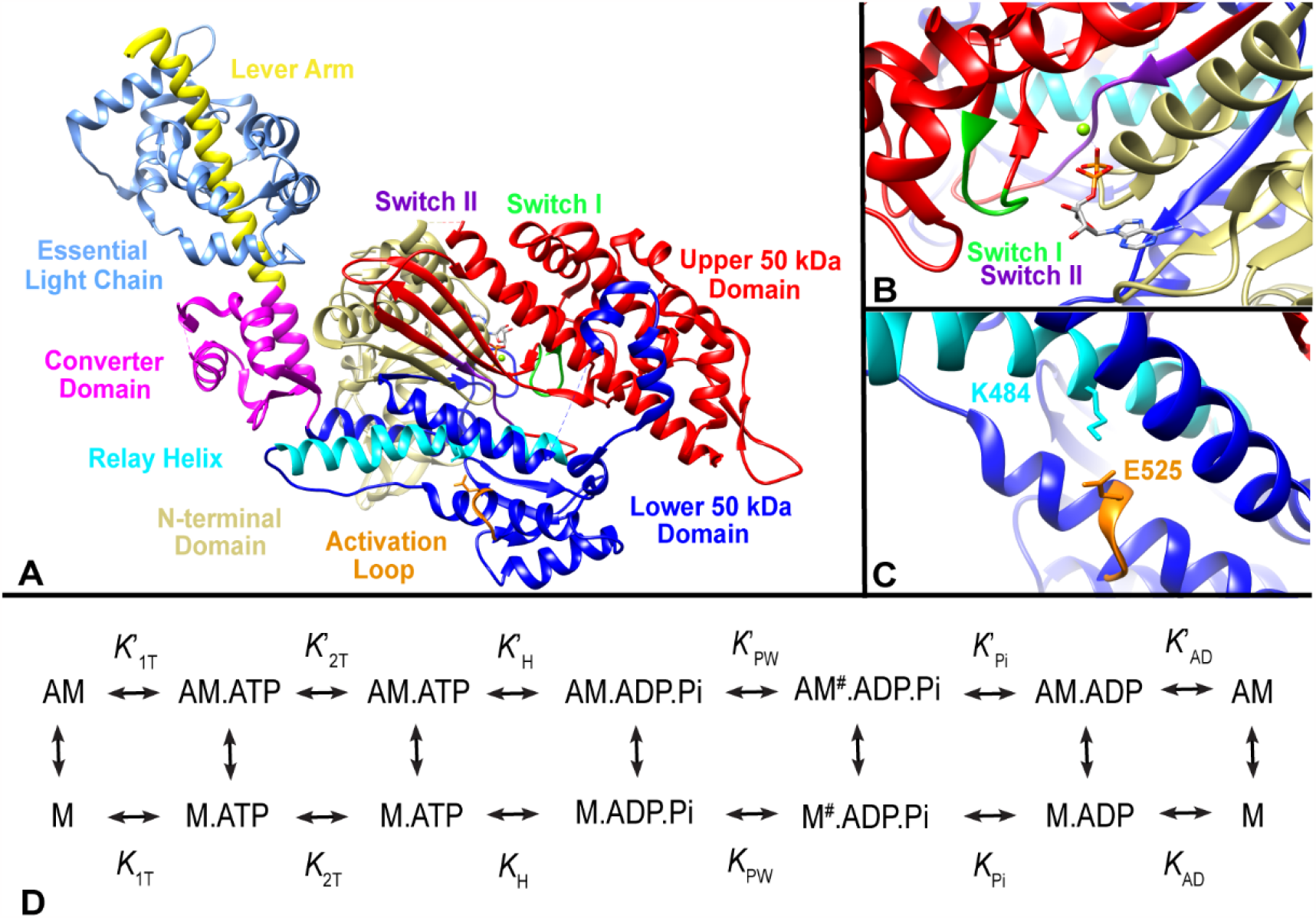
Structure of M2β S1 highlighting the E525 residue and the actomyosin ATPase cycle. **(A)** The key domains of myosin, location of E525 at the base of the activation loop within the lower 50 kDa domain, and other structural elements (e.g. switch I and switch II) of the cardiac myosin motor are highlighted (PDB 5N69). **(B)** The active site demonstrating the role of switch I and II in coordinating the gamma phosphate of ATP. **(C)** The E525 residue and its proximity to the K484 residue on the relay helix. **(D)** The kinetic scheme used to examine the key steps in the actomyosin ATPase cycle.

Cardiomyopathy is a disease of the heart muscle that results in impaired pump function and frequently leads to heart failure. The two most prevalent genetically induced cardiomyopathies, hypertrophic cardiomyopathy (HCM) and dilated cardiomyopathy (DCM), have been linked to mutations in human beta-cardiac myosin, encoded by *MYH7* (12). HCM presents clinically as an increase in hypertrophy and ventricle wall thickening, often causing diastolic dysfunction, myofilament disarray and fibrosis. In contrast, DCM, the second most common cause of heart failure, is characterized by wall thinning, left ventricular dilation, and a dramatic decrease in ejection fraction. Previous HCM and DCM cardiomyopathy *MYH7* variant studies have been successful in contributing to our understanding of how impaired cardiac myosin impacts disease pathogenesis. These studies have resulted in FDA approved small molecule therapeutics, such as Mavacamten and Omecamtiv Mecarbil (13-17). However, the complex ramifications of myosin mutations on motor function are not entirely understood. For example, the leading hypothesis suggests that beta-cardiac myosin mutations that are “*gain of function*” are associated with HCM whereas “*loss of function*” are associated with DCM (18). However, exceptions have been identified, leaving room for debate within the field (10, 19, 20).

One intriguing feature of the myosin ATPase mechanism is that actin binding dramatically accelerates ATPase activity, in some cases by several orders of magnitude (21, 22). This feature requires a precise structural mechanism that allosterically connects the actin binding region with the active site. The conserved myosin motor domain is composed of 4 subdomains, the upper and lower 50 kDa domains, the N-terminal domain, and the converter/lever arm domain (**Fig. 1A**) (2). In the current paper we focus on a region of the lower 50 kDa domain, known as the activation loop, which is thought to be crucial for triggering actin activation of ATPase activity (**Fig. 1B & C**). Previously, we studied the DCM mutation (E525K) located in the conserved activation loop (**Fig. S1**) and found that it dramatically enhanced actin-activated ATPase activity and the apparent affinity for actin (10). In addition, we found that this mutation dramatically stabilized the autoinhibited IHM and the SRX states even at physiological ionic strength (10, 23). We hypothesized that this region of myosin was crucial for stabilizing the electrostatic interactions that form the IHM, thus regulating the number of active heads in the thick filament. In the present study, we examined the impact of the E525K mutation on the transient kinetic steps of the ATPase cycle which allowed us to precisely determine the ATPase mechanism and the fraction of the ATPase cycle where myosin is bound to actin (duty ratio). We measured the power stroke rate constant using a FRET sensor of myosin lever arm rotation and performed Gaussian accelerated molecular dynamics simulations to predict structural changes and allosteric pathways. In addition, we investigated the impact of the mutation on the motile and force generating properties of S1 using unloaded and loaded motility assays. We demonstrate that the mutation uncovers a highly conserved allosteric pathway that is crucial for actin activation of the power stroke and phosphate release. Thus, our work highlights a conserved actin binding motif that can either interact with subfragment 2 (S2) to form the IHM or interact with actin to produce force, making it a major determinant of whether myosin can form an autoinhibited or force-generating myosin head.

## Results

### Steady-state actin-activated ATPase activity

In our previous work, we examined the actin-activated ATPase activity of WT and E525K M2β S1 in MOPS 20 buffer (10 mM MOPS at pH 7.0, 20 mM KCl, 1 mM EGTA, 1 mM MgCl_2_, 1 mM DTT) (10). Since previous work has demonstrated the myosin ATPase reaction is sensitive to Mg^2+^ concentration (24), we sought to use a free Mg^2+^ concentration that was saturating for the current study. Therefore, the kinetic experiments were performed in MOPS 5/4 buffer (10 mM MOPS at pH 7.0, 5 mM KCl, 1 mM EGTA, 4 mM MgCl_2_, 1 mM DTT). The actin-activated ATPase activity of M2β S1 WT and E525K was examined in three independent protein preparations and the average ATPase, at each actin concentration, was plotted as a function of actin concentration and fit to the Michaelis-Menten equation to determine the maximum rate of steady-state actin-activated ATPase activity (*k*_cat_) and the concentration where ATPase activity is one-half maximal (*K*_ATPase_) (**Fig. 2A**) (25). Our results reveal a 2.8-fold increase in the E525K *k*_cat_ (10.41 ± 0.41 s^-1^) and a 6-fold decrease in E525K *K*_ATPase_ (3.88 ± 0.77 µM) when compared to the *k*_cat_ and *K*_ATPase_ values of our WT construct, respectively (3.71 ± 0.54 s^-1^; 23.68 ± 8.14 µM) (**Table 1**). We also determined that the ATPase activity without actin, *v*_*0*_, was enhanced two-fold by the mutation (WT, 0.02 ± 0.01 s^-1^; E525K, 0.04 ± 0.01s^-1^) (**Table 1**). Single ATP turnover measurements were also utilized to examine the ATPase activity without actin (**Fig. 2B**). Myosin was first incubated with mant-labeled ATP and subsequently mixed with a high concentration (1 mM) of dark ATP. We found that the slow phase representing the SRX state was a small fraction of the fluorescence transients in both WT and E525K (∼5%) (**Table 1**). The rate constant for the fast phase, referred to as the disordered relaxed (DRX) state, was 2-fold faster in E525K compared to WT M2β S1 (**Table 1**).

**Table 1.**
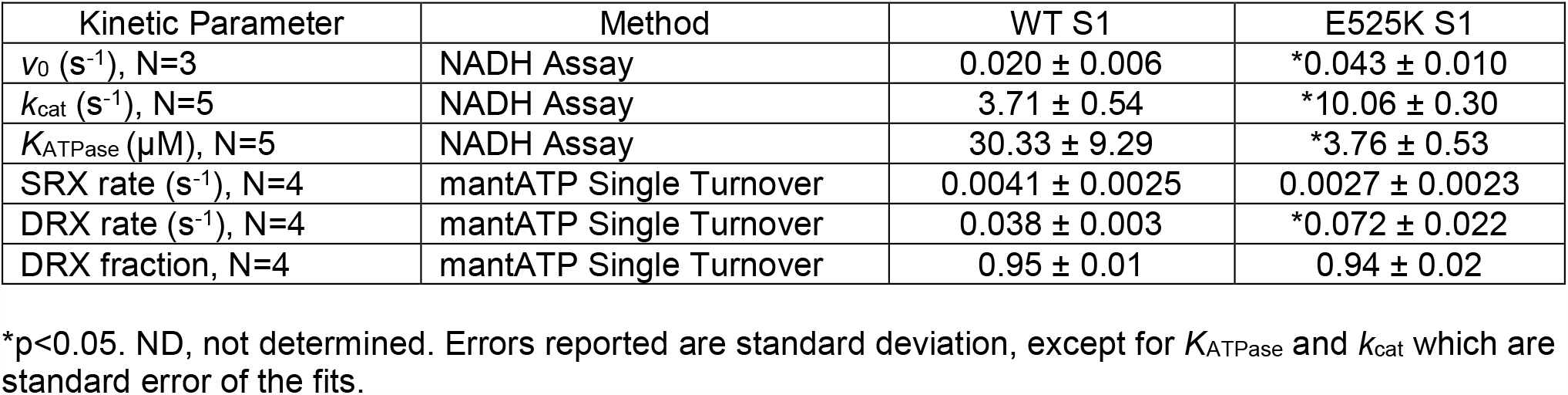
Summary of steady-state kinetic measurements.

| Kinetic Parameter | Method | WT S1 | E525K S1 |
| --- | --- | --- | --- |
| $v_0$ ($s^{-1}$), N=3 | NADH Assay | $0.020 \pm 0.006$ | *$0.043 \pm 0.010$ |
| $k_{cat}$ ($s^{-1}$), N=5 | NADH Assay | $3.71 \pm 0.54$ | *$10.06 \pm 0.30$ |
| $K_{ATPase}$ ($\mu M$), N=5 | NADH Assay | $30.33 \pm 9.29$ | *$3.76 \pm 0.53$ |
| SRX rate ($s^{-1}$), N=4 | mantATP Single Turnover | $0.0041 \pm 0.0025$ | $0.0027 \pm 0.0023$ |
| DRX rate ($s^{-1}$), N=4 | mantATP Single Turnover | $0.038 \pm 0.003$ | *$0.072 \pm 0.022$ |
| DRX fraction, N=4 | mantATP Single Turnover | $0.95 \pm 0.01$ | $0.94 \pm 0.02$ |

\* $p < 0.05$ . ND, not determined. Errors reported are standard deviation, except for $K_{ATPase}$ and $k_{cat}$ which are standard error of the fits.

**Figure 2.**
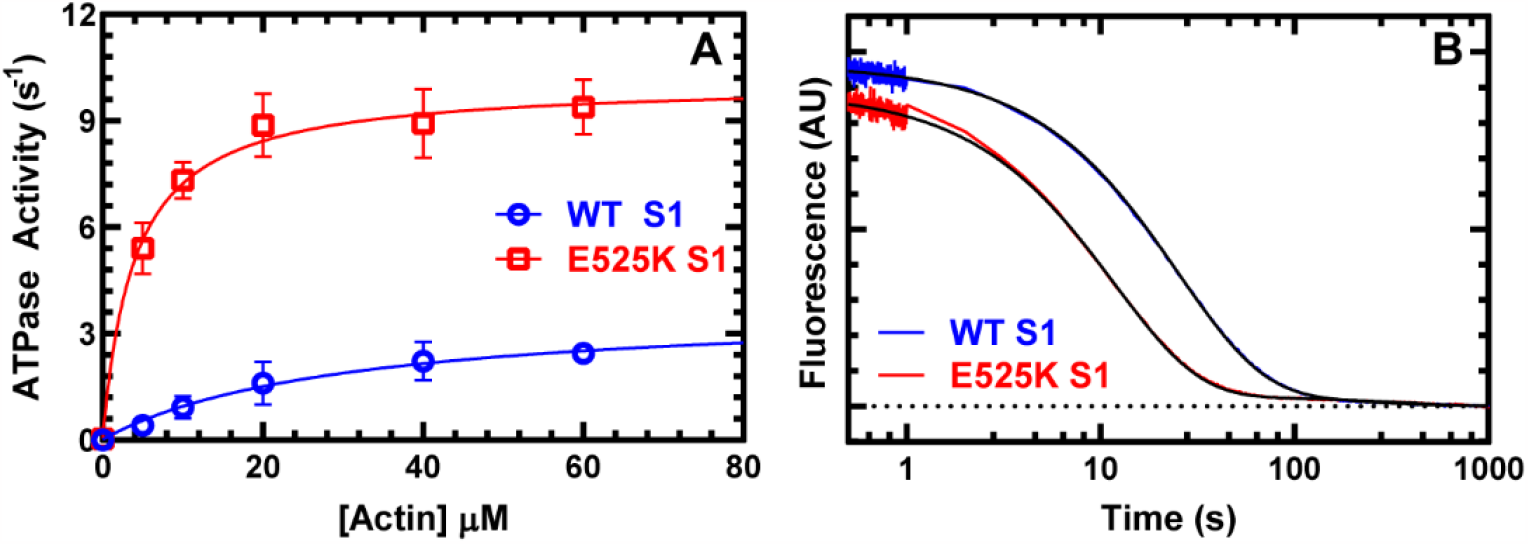
Steady-state ATPase kinetics. **(A)** The actin-activated ATPase activity of purified M2β S1 WT and E525K was determined as a function of actin concentration. The data points represent the average ± SD from three protein preparations. **(B)** Turnover of mant-ATP by M2β S1 WT and E525K was examined for 1000 s after (0.25 µM) myosin was pre-incubated with mant-ATP (1 µM) for ∼30 s and then mixed with saturating unlabeled ATP (2 mM). Fluorescence transients were best fit to a double exponential function.

### Impact of E525K on transient kinetic steps of the ATPase cycle

We investigated the impact of the E525K mutation on the predominant steps of the actomyosin ATPase cycle. The results were interpreted in the context of a simplified kinetic scheme of the actomyosin ATPase cycle (**Fig. 1D**) (11, 26). The equilibrium constants in this scheme, each with associated forward (right) and reverse (left) rate constants (not shown for simplicity), are referred to throughout the manuscript.

The ATP binding and hydrolysis steps were measured by monitoring the tryptophan fluorescence enhancement that occurs when M2β S1 (1 μM) was mixed with varying concentrations of ATP. Fluorescence transients were best fit to a double exponential function and the rate constants were plotted as a function of ATP concentration (**Fig. S2A & C**). The fast phase rate constants were dependent on ATP concentration and were fit to a hyperbolic function to determine the maximum rate of ATP hydrolysis, *k*_+H_ + *k*_-H_ (s^-1^), and dependence on ATP concentration, *K*_0.5_ (µM) (**Fig. S2B**). E525K showed a slightly reduced maximum rate of ATP hydrolysis (100.4 ± 3.2 s^-1^) compared to WT (127.9 ± 4.2 s^-1^) while *K*_0.5_ values were similar for both constructs (WT, 21.6 ± 2.7 µM; E525K, 20.9 ± 2.6 µM) (**Table 2**). The second-order rate constant for ATP binding to myosin, *K*_*1T*_*k*_*+*2T_ (µM^-1^s^-1^), was determined by the linear dependence of the fast phase on ATP concentration at lower concentrations of ATP, which was similar for both constructs (WT, 3.5 ± 0.2 µM^-1^·s^-1^; E525K, 2.7 ± 0.2 µM^-1^·s^-1^) (**Table 2**). The rate constant of the slow phase, which represented 10-20% of the fluorescence signal, was independent of ATP concentration and similar in WT and E525K (∼20 s^-1^).

**Table 2.** Summary of transient kinetic measurements.

| Kinetic Step | Method | Rate/equilibrium constant | WT S1 | E525K S1 |
| --- | --- | --- | --- | --- |
| ATP binding to M<br>N=3 | Tryp. Fluor | $K_{1T}k_{2T}$ ($\mu\text{M}^{-1}\text{s}^{-1}$)<br>$K_{0.5}$ ($\mu\text{M}$) | $3.5 \pm 0.2$<br>$21.6 \pm 2.7$ | *$2.7 \pm 0.2$<br>$20.9 \pm 2.6$ |
| ATP hydrolysis<br>N=3 | Tryp. Fluor | $k_{+H} + k_{-H}$ ($\text{s}^{-1}$) | $127.9 \pm 4.2$ | *$100.4 \pm 3.2$ |
| ATP binding to AM<br>N=5 | Pyrene Actin | $K'_{1T}k'_{+2T}$ ($\mu\text{M}^{-1}\text{s}^{-1}$) | $6.8 \pm 0.5$ | $8.0 \pm 0.8$ |
| Dissociation of AM by ATP<br>N=4 | Pyrene Actin | $k'_{+2T}$ ($\text{s}^{-1}$)<br>$K_{0.5}$ ($\mu\text{M}$) | $827.8 \pm 111.8$<br>$99.8 \pm 22.6$ | *$1993 \pm 212.4$<br>$212.4 \pm 99.1$ |
| Power Stroke (fast)<br>@ 30 $\mu\text{M}$ actin, N=3 | FRET | $k'_{+PW}$ ($\text{s}^{-1}$)<br>$K_{0.5}$ ($\mu\text{M}$) | $10.7 \pm 6.8$<br>ND | *$43.7 \pm 8.0$<br>$17.4 \pm 9.8$ |
| Power Stroke (slow)<br>@ 30 $\mu\text{M}$ actin, N=3 | FRET | $k'_{+PWS}$ ($\text{s}^{-1}$) | $1.0 \pm 0.3$ | $2.3 \pm 1.7$ |
| Pi release (fast),<br>@ 30 $\mu\text{M}$ actin, N=1-2 | MDCC-PBP | $k'_{+Pi}$ ($\text{s}^{-1}$)<br>$K_{0.5}$ ($\mu\text{M}$) | $22.4 \pm 4.0$<br>ND | $65.2 \pm 1.1$<br>$8.9 \pm 3.6$ |
| Pi release (slow)<br>N=2 | MDCC-PBP | $k'_{+PiS}$ ($\text{s}^{-1}$)<br>$K_{0.5}$ ($\mu\text{M}$) | $2.6 \pm 0.3$<br>$61.1 \pm 58.3$ | $2.5 \pm 0.1$<br>$1.3 \pm 1.1$ |
| ADP dissociation from AM<br>N=4-7 | Pyrene Actin<br>mantADP | $k'_{+ADP}$ ($\text{s}^{-1}$)<br>$k'_{+ADP}$ ($\text{s}^{-1}$) | $116.5 \pm 16.0$<br>$291.4 \pm 27.0$ | $114.5 \pm 12.1$<br>*$327.9 \pm 20.2$ |

\* $p < 0.05$ . ND, not determined. Errors reported are standard error of the fit, except for Power stroke, Pi-release, and ADP dissociation which are standard deviation.

ATP-induced dissociation from actin was measured by mixing M2β S1 (0.5 μM) complexed with equal molar pyrene labeled actin (0.5 μM) with increasing concentrations of ATP. Fluorescence transients were best fit to a single exponential function and the rate constants were fit to a hyperbolic function to determine the maximum rate of transition into the weakly bound states, *k’*_*+*2T_ (s^-1^), and the equilibrium constant for the initial interaction of ATP with actomyosin (*K*’_1T_) (*k*_obs_ = [ATP]*k*’_+2T_/([ATP]+*K*_0.5_), where *K*_0.5_ = 1/*K*’_1T_) (**Fig S3A & C**). The E525K mutation enhanced *k’*_*+*_2T (1993 ± 212.4 s^-1^) 2.4-fold compared to WT (827.8 ± 111.8 s^-1^) (**Table 2**). The second-order rate constant, *K’*_*1T*_*k’*_*+2T*_ (µM^-1^s^-1^), determined from the linear fit of the fast phase at lower ATP concentrations, was similar in both constructs (**Fig S3B**). The *K*_0.5_ was not significantly different between constructs (**Table 2**).

A sequential mix stopped-flow configuration was used to measure the rate of actin-activated phosphate release. Actin-activated phosphate release was measured by mixing M2β S1 with sub stoichiometric ATP and aging for 10 seconds to allow ATP binding and hydrolysis followed by a second mix with actin in the presence of phosphate-binding protein (final concentrations: 1uM M2β, 0.9 µM ATP, 4 µM PBP, and 5-30 μM actin). Phosphate release fluorescent transients collected under single turnover conditions were best fit by a double exponential function (**Fig. 3**). The slow phase, *k’*_*+*Pi_ (slow) (s^-1^), was similar for both the WT and mutant constructs at 30 μM actin (WT, 2.6 ± 0.3 s^-1^; E525K, 2.9 ± 0.4 s^-1^) (**Fig. 3B, Table 2**). The mutant does demonstrate an enhanced actin dependence of *k’*_*+*Pi_ (slow), as it is able to plateau at a much lower actin concentration. The E525K construct produced a 3.4-fold increase in the fast phase of actin-activated phosphate release, *k’*_*+*Pi_ (fast) (s^-1^), compared to WT at 30 μM actin (WT, 22.4 ± 4.0 s^-1^; E525K, 65.2 ± 1.1 s^-1^) (**Fig. 3A, Table 2**). The maximum rate of *k’*_*+*Pi_ (fast) was found to be 76.9 ± 11.9 for the mutant but was not determined for WT. E525K demonstrated an increased relative amplitude of the fast phase while WT was dominated by the slow phase (fast phase relative amplitude; WT, 9.2 ± 3.0 %; E525K, 48 ± 17 %).

**Figure 3.**
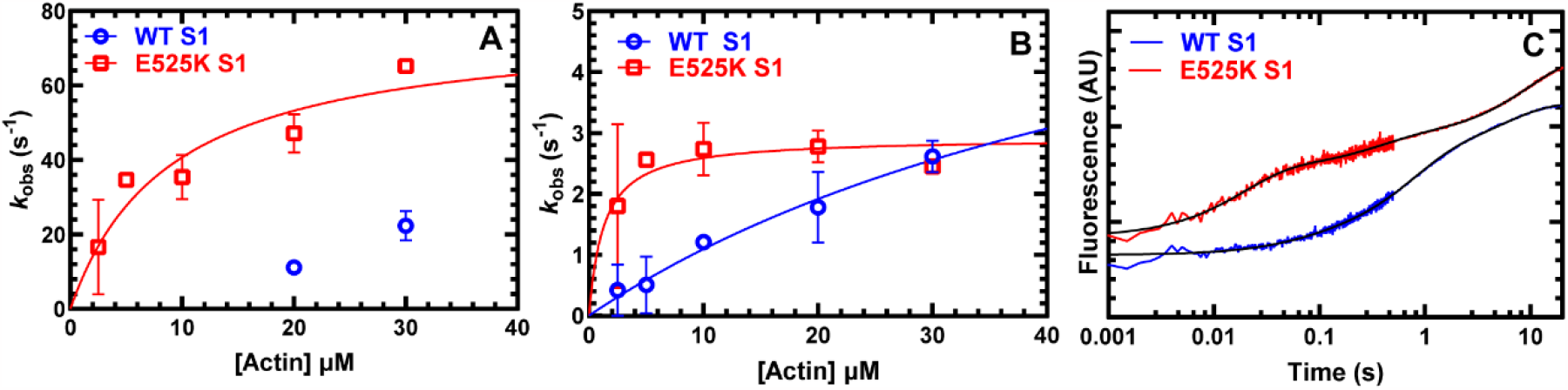
Actin-activated phosphate release. The phosphate binding protein (MDCC-PBP) was used to monitor the phosphate release step using sequential-mix experiments. M2β S1 was mixed with substoichiometric ATP, aged for 10 s, and then mixed with actin and MDCC-PBP (final concentrations: 1 µM M2B S1, 0.9 µM ATP, 5 µM MDCC-PBP, and 5-30 μM actin). The rate constants of the **(A)** fast and **(B)** slow phases were plotted as a function of actin concentration and fitted to a hyperbolic function. **(C)** Representative fluorescence transients in the presence of 20 μM actin (average of two transients) are shown fitted to a double exponential function. Data points at each actin concentration represent the average ± SD of two experiments from two separate protein preparations.

We examined the rate constant for ADP release from actomyosin, *k’*_*+*ADP_ (s^-1^), by mixing a pyrene actomyosin-ADP complex (0.5 µM actomyosin: 10 µM ADP) with saturating ATP (2 mM) (**Fig. S4A**). Similar rate constants were observed for WT and E525K (WT, 122.9 ± 11.9 s^-1^; E525K, 118.9 ± 10.2 s^-1^) (**Table 2**). The rate constant for ADP release from actomyosin was also calculated using a second method with mant-ADP (**Fig. S4B**). The actomyosin mant-ADP complex (0.5 µM actomyosin: 10 µM mant-ADP) was mixed with saturated (1 mM) ATP. The rate constants were faster than with unlabeled ADP, as we reported previously (27, 28), and we observed a slight increase with the mutant (WT, 291.4 ± 27.0 s^-1^; E525K, 327.9 ± 20.2 s^-1^) (**Table 2**).

In a previous study we employed a FRET approach for measuring the rotation of the lever arm in M2β S1 by monitoring the transition from the pre- to post-power stroke states (28). Power stroke FRET was examined by exchanging Alexa488 labeled RLC onto M2β S1 (donor) and then utilizing Cy3-labeled nucleotides bound to the active site as the acceptor. To measure the actin-activated power stroke, M2β S1 A488RLC was mixed with substoichiometic Cy3ATP, aged for 10 seconds to allow for hydrolysis to occur, and then mixed with actin (final concentrations 0.5 µM M2B, 0.45 µM Cy3ATP, and 5-30 μM actin). The traces were best fit to a double exponential function, especially at higher actin concentrations (10, 20, and 30 µM) (**Fig. 4**). The fast phase of the power stroke measurements was hyperbolically dependent on actin concentration in E525K, which allowed us to determine the maximum rate of the power stroke (*k*_+PWS_ = 66.9 ± 18). In WT M2β S1 we also observed two exponentials in the power stroke transients, but the fast phase rate constant was linearly dependent on actin concentration preventing us from determining the maximum rate. Thus, we report the power stroke rate constant at 30 µM actin to compare the WT and mutant constructs. The E525K mutation increased the fast phase rate, *k’*_PW_ (fast) (s^-1^), 4-fold (WT, 10.7 ± 6.8 s^-1^; E525K, 43.7 ± 8.0 s^-1^) while the slow phase rate, *k’*_PW_ (slow) (s^-1^) was not significantly different (WT, 1.0 ± 0.3 s^-1^; E525K, 2.3 ± 1.7 s^-1^) at 30 µM actin (**Table 2**). The relative amplitude of the fast phase of the fluorescence transients at 30 µM actin was larger in E525K than WT (89 ± 0.01% and 53 ± 0.16%, respectively)

**Figure 4.**
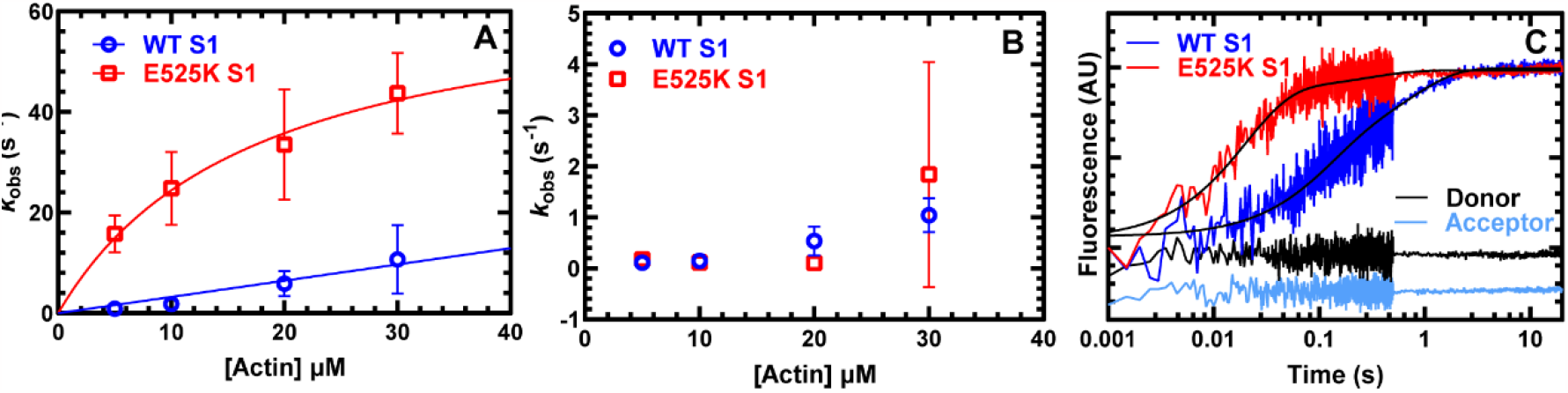
Actin-activated power stroke. Actin-activated power stroke rate constants were measured by monitoring the fluorescence enhancement of Alexa 488 during actin-activated product release. Sequential-mix stopped-flow experiments were performed by mixing M2β S1 A488RLC Cy3ATP, aged for 10 s for hydrolysis to occur, and then mixed with actin (final concentrations: 0.5 µM M2B S1, 0.45 µM ATP, and 5-30 μM actin). The fluorescence transients were best fit by a double exponential function. **(A)** The rate constants of the fast phase were plotted as a function of actin concentration and fitted to a hyperbolic function for E525K and WT. **(B)** The slow phase of the power stroke was relatively independent of actin concentration. **(C)** Representative fluorescence transients in the presence of 30 μM actin (average of two transients) are shown fitted to a double exponential function. The donor-only and acceptor-only controls are shown for comparison. Data points at each actin concentration represent the average ± SD of three experiments from separate protein preparations.

### Kinetic simulations and duty ratio

In order to determine the predominate kinetic pathway of WT and E525K S1 we performed kinetic simulations using the individual rate constants measured, as well as taking into consideration other published studies (**Table S1**). We favor a model in which myosin proceeds through actin-bound (*K*’_H_) and actin-detached (*K*_H_) ATP hydrolysis pathways. In addition, the actin-activated power stroke (*K*’_PW_) occurs prior to the release of phosphate (*K*’_Pi_). The rate constants for the kinetics steps not measured in the current study were adjusted manually until the simulated data fit best to the experimental data. This model fits reasonably well to the steady-state ATPase (**Fig. S5A**) as well as the phosphate release (**Fig. S5B & C**) and power stroke (**Fig. S5D & E**) rate constants for both the mutant and WT.

The duty ratio at 30 µM actin was estimated using the following equation where *k*_H_ is the maximum rate of ATP hydrolysis measured by intrinsic tryptophan fluorescence, *k’*_PW_ is the fast phase of the actin-activated power stroke rate constant, and *k’*_D_ is the ADP release rate constant monitored with pyrene actin fluorescence.

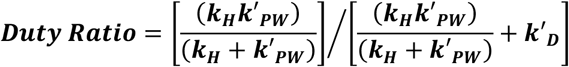

Using this equation, we calculate that the mutant enhances the duty ratio approximately 3-fold (WT, 0.08; E525K, 0.21).

### Impact of E525K on actin sliding velocity and contractile force

We performed both unloaded and loaded *in vitro* actin gliding assays with WT and E525K M2β S1. All motility assays were performed at room temperature (∼22-23°C) in motility buffer (10 mM MOPS, pH 7.0, 50 mM KCl, 1 mM EGTA, 1 mM MgCl_2_, 1 mM DTT), which we found was optimal for performing both loaded and unloaded motility. The unloaded sliding velocity was about 20% faster in the E525K mutant, and this increase was consistent over a wide range of densities. We fit the density dependent data to a hyperbolic function to determine the maximum velocity (*V*_MAX_) and density at which the velocity is one-half maximal (*K*_0.5_) (**Fig. 5; Table 3**). The *V*_MAX_ was 20% faster with the mutant and the *K*_0.5_ was shifted two-fold lower, consistent with an increase in duty ratio. We also performed *in vitro* motility in the presence of varied concentrations of alpha-actinin to introduce a load to the assay. We found that E525K required significantly more alpha-actinin to slow the *in vitro* gliding velocities 50% (EC50=16.2±1.0 and 10.6±1.1 for E525K and WT, respectively) when the data in Figure 5B were fit to a standard dose response curve. We utilized the assumptions from Greenberg and Moore to convert alpha-actinin concentration to viscous load, which allowed us to fit the data to the force-velocity relationship (29). We determined that the mutant generated a larger (68%) maximum ensemble force (or isometric force). We also plotted the force as a function of power and found that E525K generated a 2-fold higher peak power and the force at which peak power was observed was also increased (68%). Our results strongly complement our kinetic characterization and clearly demonstrate that this mutation enhances both enzymatic and motor properties.

**Table 3.** Summary of *in vitro* motility measurements.

| **Motor Property** | **WT S1** | **E525K S1** |
| --- | --- | --- |
| Maximum unloaded sliding velocity ($V_{MAX}$) ($\text{nm}^{-1}\text{s}^{-1}$) | $3663 \pm 63$ | *$4535 \pm 44$ |
| Density at which unloaded $V$ is $\frac{1}{2} V_{MAX}$ ($\mu\text{M}$) | $0.239 \pm 0.010$ | *$0.121 \pm 0.004$ |
| EC50 for alpha-actinin ($\mu\text{g/mL}$) | $10.6 \pm 1.1$ | *$16.2 \pm 1.0$ |
| Maximum ensemble force ($F_{MAX}$) (pN) | $796 \pm 51$ | *$2040 \pm 79$ |
| Peak power (fW) | 0.39 | 0.81 |
| Force at which peak power occurs (pN) | 231 | 388 |

\* $p < 0.05$ . ND, not determined. Errors reported are standard error of the fit (N=3).

**Figure 5.**
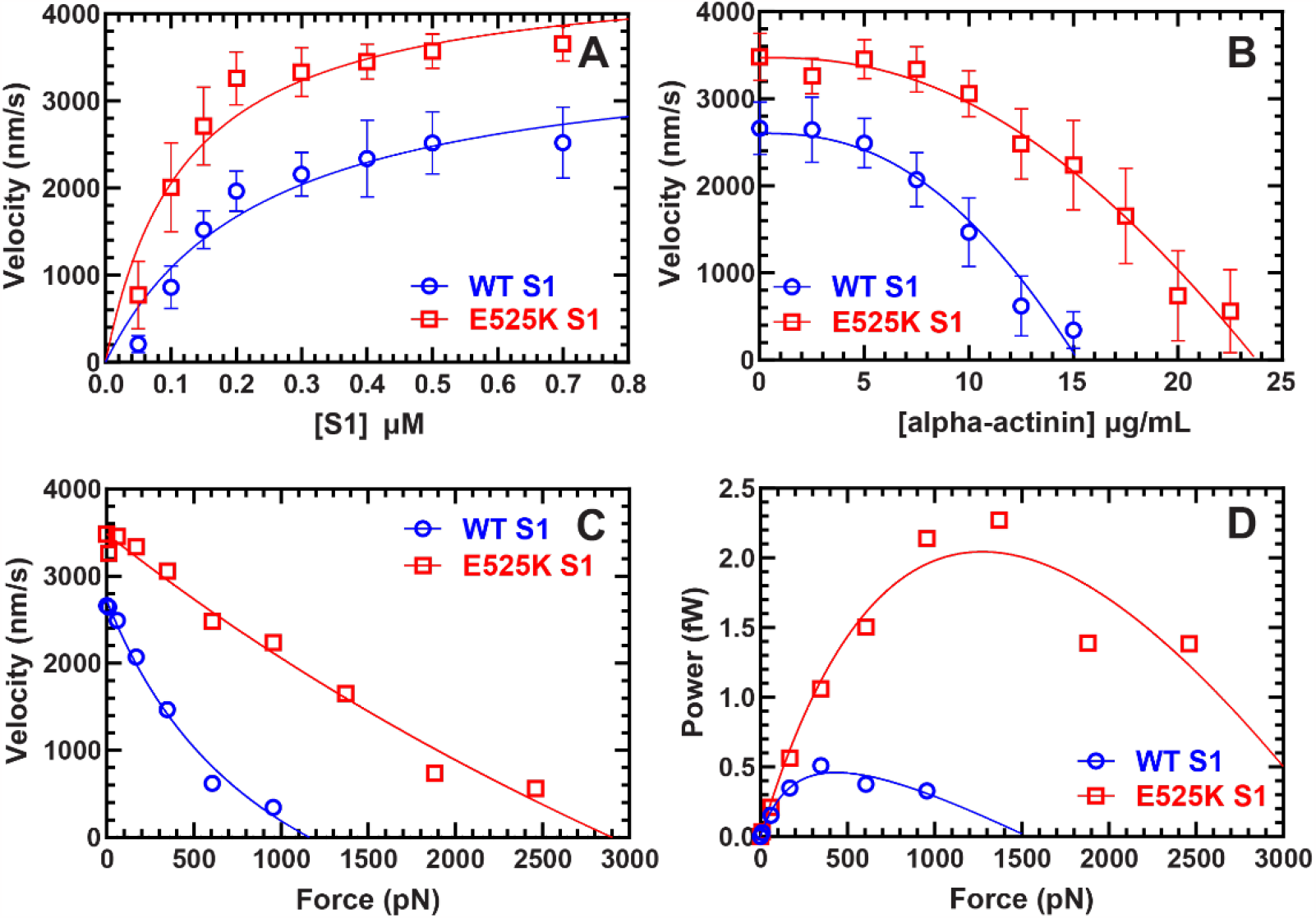
Actin gliding, ensemble force, and power. **(A)** Actin sliding velocities were quantified across a range of myosin surface densities in an *in vitro* motility assay. Surface density modulation was achieved by varying myosin incubation concentrations within the range of 0.05-0.7 μM. Data points were presented as the mean ± SD, with measurements conducted using three separate protein preparations. A total of 150 actin filaments (50 actin filaments per protein preparation, N = 3) were assessed for each condition. **(B)** To investigate the impact of frictional load, alpha-actinin, an actin-binding protein was employed. The total concentration of M2β S1 loaded onto the surface remained constant at 0.4 μM, while the concentration of alpha-actinin was varied within the range of 0 - 22.5 μg/mL until a point of actin filament movement cessation was reached. A total of 100 actin filaments from three distinct protein preparations were subjected to testing. Data were fitted to the elastic model (29). The error bars presented are the mean ± SD. **(C)** Using the data from **Fig.5B**, alpha-actin in concentrations were converted to viscous load, which allowed us to fit the data to the force-velocity relationship. The data were fit to the Hill equation as described by Greenberg and Moore (29), resulting in the determination of the maximum ensemble force (*F*_MAX_) **(D)** Data from force-velocity relationship (**Fig. 5C**) were transformed into power by multiplying the force by the velocity, subsequently facilitating the generation of power-force plots. Peak power values and the corresponding force at which peak power occurred were determined by plotting and fitting the curve to the Hill equation. These results are summarized in **Table 3**.

### Atomistic simulations reveal mechanistic differences between WT and E525K actomyosin

To investigate the impact of the E525K mutant on the myosin power stroke at an atomistic level, we carried out Gaussian accelerated MD (GaMD) simulations on the pre-power stroke (PPS) actomyosin (**Fig. 6A**) (30). The E525K simulations were set up based on models obtained for the WT PPS actomyosin complex (31). Our analysis shows that the WT PPS myosin can form a stable salt bridge between E525 and K484, which has a most probable distance of 3.5Å (cyan curve in **Fig. 6C**). In the case of E525K simulations, this salt bridge is destabilized and the most probable K525-K484 distance increases to 6.5 Å (orange curve in **Fig. 6C**), accompanied by rotations of the relay helix (RH, **Fig. 6B**). The E525K mutant displays more pronounced variations in the relay helix (RH) motion compared to the WT case, as shown in **Fig. S6**. These changes further induce a metastable state of the active site, characterized by a widened gap between switch I and II (**Fig. 6D**). The free energy profile along the distance between switch I and switch II finds this metastable state (17Å) to be exclusive to the E525K simulations (highlighted by the black dotted line in **Fig. 6E**). Given that switch I and II are proposed to form the “back door” for phosphate release, the widening of the gate represents an intermediate state potentially facilitating phosphate release. Our results suggest an allosteric mechanism that is adopted by the E525K mutant to promote Pi release and power stroke transition (further discussed below).

**Figure 6.**
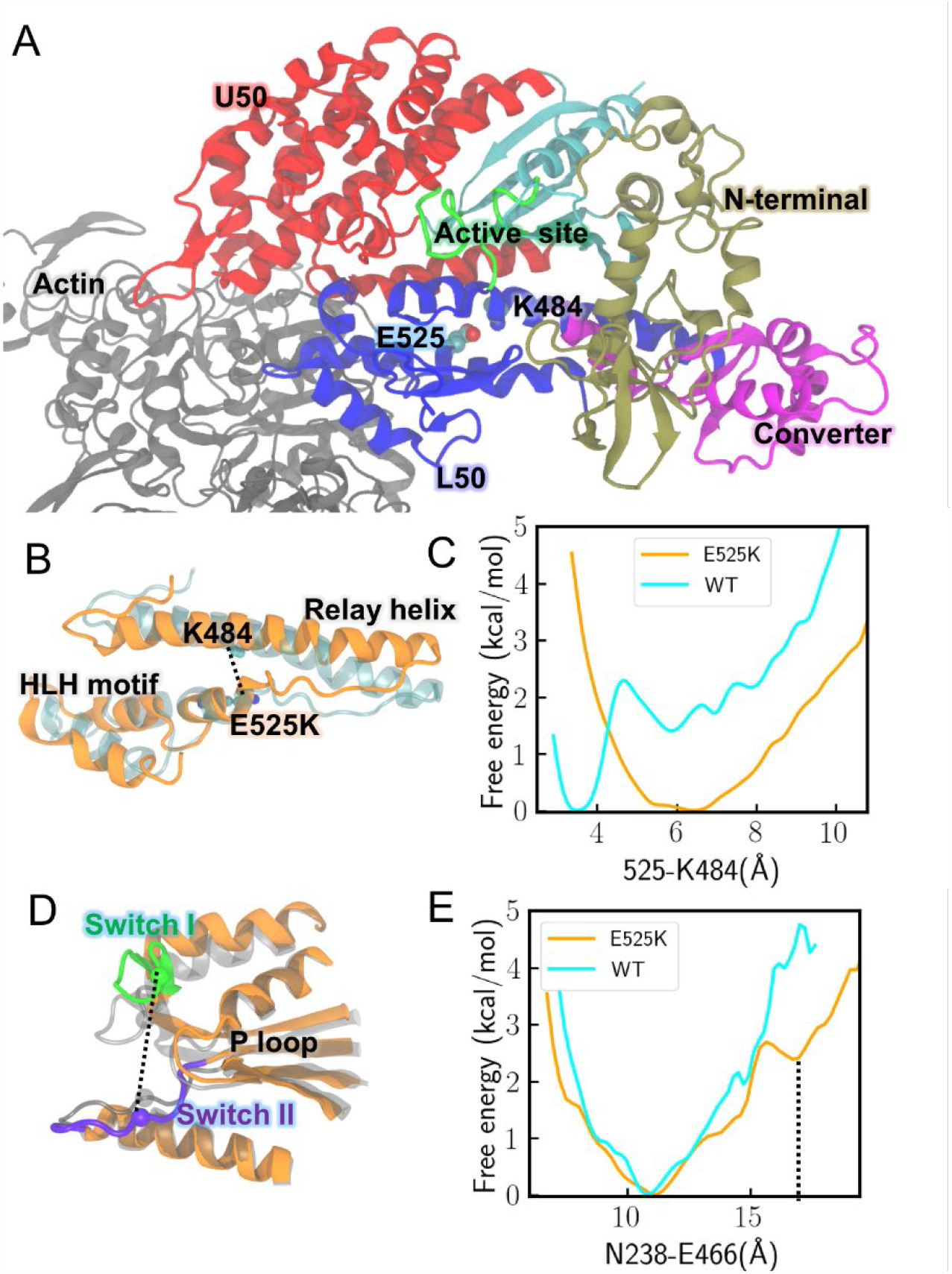
GaMD simulations reveal differences between WT and E525K myosin. **(A)** A structural model for the pre-power stroke (PPS) actomyosin with domains highlighted in different colors. **(B)** Representative conformations of the relay helix and helix loop helix (HLH) motif for the E525K actomyosin simulations (orange) and WT actomyosin simulations (cyan) at the PPS state. **(C)** The free energy profiles projected along the distance between residues 525 and K484. **(D)** For the E525K mutant, myosin active site at the most probable state is shown in transparent gray, whereas the active site at a metastable state is shown in orange (switch I in green, switch II in purple). **(E)** The free energy profiles projected along the distance between N238 (switch I) and E466 (switch II). The metastable state shown in panel D is highlighted by the black dotted line. This state, which is only observed in the E525K simulations (orange curve), likely facilitates Pi release.

## Discussion

We completed a comprehensive analysis of the E525K mutation in monomeric M2β S1, which allowed us to determine the impact of the mutation on the ATPase mechanism, duty ratio, motile properties, and structural mechanism of force generation. We revealed that the 3-fold increase in steady-state actin-activated ATPase activity is explained primarily by significantly enhanced actin-activated power stroke and phosphate release rate constants. With only minor changes to the other steps in the ATPase cycle, the enhancement in the power stroke/phosphate release increases the duty ratio approximately 3-fold in the E525K mutation. Our *in vitro* motility results align with the kinetic characterization by demonstrating increases in actin gliding velocity, ensemble force, and power. We also performed Gaussian molecular dynamics simulations, which allowed us to propose a specific allosteric pathway for actin activation of the power stroke and phosphate release. Lysine 484 in the relay helix forms an essential salt bridge with glutamate 525 in the activation loop that is abolished by the mutation. The resulting increased flexibility of the relay helix and activation loop facilitates the enhanced actin activation mechanism. In a previous study, we demonstrated that the E525K mutation dramatically stabilizes the autoinhibited state in dimeric heavy meromyosin (HMM). Therefore, we propose that an important feature of the autoinhibited state is its ability to sequester the lower 50 kDa domain region containing the activation loop from interacting with actin, subsequently preventing initiation of the highly conserved actin activation pathway. In addition, this study sheds light on the balance between intrinsic motor properties and autoinhibition, and the interplay between these factors that ultimately leads to the reduced force and power observed in DCM patients.

### Impact of DCM mutant E525K on intrinsic ATPase cycle kinetics

Steady-state ATPase measurements uncovered that the E525K mutation increased the actin-activated ATPase approximately 3-fold and dramatically reduced (6-fold) the actin concentration required to reach one-half maximal ATPase. Impressively, the catalytic efficiency (*k*_cat_/*K*_ATPase_) was improved 17-fold in the mutant. We did observe a 2-fold increase in ATPase activity in the absence of actin, demonstrating the mutation alters the rate-limiting step, likely phosphate release, in the absence of actin. The current study differs from our previous work, in that we used a buffer with lower ionic strength and a higher magnesium concentration (MOPS 5/4). Transient kinetic analysis revealed the mutation only resulted in minor changes to ATP binding, hydrolysis, and ATP-induced dissociation from actin, with no significant changes or minor changes in ADP release. Actin-activated phosphate release experiments, measured using the phosphate binding protein, revealed a 3-fold increase in the fast phase of the fluorescence transients, similar to that observed with steady-state ATPase. However, the slow phase was quite similar in both mutant and WT. In general, our phosphate release results agree with what has been previously reported in the literature for WT M2β S1. *Tang et al*. (2019) reported two phases in the phosphate release transients at 30 µM actin with similar slow and fast phase rate constants (27). However, another study from our group (*Tang et al. 2021*) revealed transients with a subtle lag followed by a single exponential phase that was equivalent to the slow phase in our current study and the previous *Tang et al. 2019* study (27, 28). Thus, M2β S1 phosphate release transients are dominated by a slow component and a minor fast component that can be difficult to detect since its relative amplitude is quite small (10-20%). Two other studies performed phosphate release with recombinant M2β HMM and regulated thin filaments and found fluorescence transients with fast and slow components with similar rate constants to our work, while the fast component contained a larger relative amplitude (32, 33).

The actin-activated power stroke was measured using a FRET approach that detects the rotation of the lever arm during the transition from the pre- and post-power stroke states. Transients were best fit to a double exponential function, with similar rate constants for WT M2β S1 as were found in our previous study (*Tang et al. 2021*) (28). FRET experiments demonstrated the mutation enhanced the fast power stroke rate constant 4-fold. Overall, the fast phase of the power stroke and phosphate release were similar in both the mutant and WT constructs, suggesting the two steps may be indistinguishable in cardiac myosin. This is in contrast to our studies on myosin V that demonstrated the power stroke occurs faster than phosphate release and provided direct evidence that the power stroke precedes the slower phosphate release step (34). The duty ratio can be calculated by utilizing the power stroke rate constant to determine the transition into the strongly bound state and the ADP release rate constant for determining exit from the strongly bound states. Using this equation, we calculate that the mutant enhances the duty ratio 3-fold.

It is tempting to speculate that the slow phase in the power stroke and phosphate release transients is derived from the actin-bound hydrolysis step. Previous work from White and colleagues has provided strong evidence that this pathway exists in the fast skeletal muscle myosin ATPase mechanism and later in porcine cardiac myosin (33, 35). Our data is best fit to a scheme that includes two pathways, an actin-bound ATP hydrolysis pathway and the canonical pathway with ATP hydrolysis of actin following by tightly coupled power stroke and phosphate release (see **Table S1**) (35-37). Other pathways cannot currently be ruled out, such as a slow and fast pathway for phosphate release, and thus further studies may be necessary to define the rate-limiting steps in the cardiac muscle myosin ATPase cycle.

### E525K enhances actin gliding, ensemble force, and power

By characterizing the motile properties of the E525K mutant, we found that the enhanced duty ratio observed in our kinetic studies translates to increased ensemble force and power. Several other studies have compared duty ratio and ensemble force and power using the alpha-actinin loaded motility assay (29, 38-40). Our results demonstrate that E525K does not disrupt the mechanism of force production, which is thought to be rotation of the lever arm while myosin is tightly bound to actin. Thus, our motility results reinforce our measurements of the power stroke rate constant, which found that lever arm rotation was accelerated by the mutation and suggest the mutation does not impair the conserved structural mechanism of force generation but simply accelerates key aspects of the mechanism. The more rapid transition into the strongly bound state with little change in the transition into the weakly bound intermediates will allow more force generating motors to interact with actin filaments at any given time and increase the ensemble force of the mutant compared to WT. Alternatively, the individual E525K mutant motors may have a greater force generating capacity. This could be accomplished by changing the mechanical properties of the lever arm, which was observed in other studies (41, 42). Thus, it will be important to examine the single molecule mechanical properties of the E525K mutation to further investigate the mechanism of how it enhances ensemble force and power.

The increase in velocity observed with the mutant was unexpected since the ADP release rate constant, which typically correlates with unloaded sliding velocity, was unchanged or only slightly changed (43). We found a 12% change in the mant-ADP release rate constant and no change in ADP release from pyrene actomyosin. Classic models of muscle contraction predict that the ADP release rate constant should control the detachment rate which suggests detachment limited sliding velocity (44). Early studies by Barany found a correlation between ATPase rate, which is thought to be attachment limited in muscle myosins, and muscle shortening velocity (43). Recent studies by the Baker and Cremo groups have developed a model that suggests maximum velocity is dependent on both attachment (40%) and detachment (60%) kinetics (45). Thus, the E525K mutation which accelerates ATPase and attachment kinetics, but causes little change in detachment kinetics, seems to provide further evidence for attachment-limited models of *in vitro* actin gliding.

### Allosteric mechanism of actin activation

Our work has uncovered key structural details of the allosteric pathway associated with actin activation of the ATPase cycle. The activation loop, located in the lower 50 kDa domain of the actin binding region, is a highly conserved loop across myosin isoforms that was first discovered in *Dictyostelium* myosin (21). The loop was termed “activation loop” for its ability to influence actin activation kinetics in *Dictyostelium* myosin without introducing any other significant changes to the ATPase mechanism in the absence of actin (21). This loop varies in size depending on the specific isoform, ranging from about 5 to 10 residues in length. Each activation loop across the myosin family contains a positively charged amino acid at the tip, such as arginine 520 in *Dictostelium* myosin and lysine 526 in cardiac myosin (see **Fig. S1** for alignment). This positive charge is thought to be essential for interactions with the N-terminus of actin, which promotes the rotation of the lower 50 kDa domain during the transition from pre- to post-power stroke states and the release of phosphate (22).

The conserved intrinsic structural transitions during myosin force generation are still being elucidated. Actin-bound myosin, with the products of ATP hydrolysis in the active site, undergoes key conformational changes that require allosteric communication between the active site and the actin binding region as well as the converter/lever arm. The changes in the active site promote the release of phosphate, while the changes in the converter/lever arm promote rotation of the lever arm into the post-power stroke state. During this process ADP remains in the active site, while phosphate has been proposed to exit though a “back door” mechanism gated by an essential salt bridge between switch I and switch II (46, 47). In cardiac myosin, glutamate 525 at the base of the activation loop forms a salt bridge with lysine 484 of the relay helix. Substituting lysine at the 525 position disrupts this salt bridge making the relay helix more flexible. Increased flexibility of the relay helix induces a metastable state in which the gap between switch I and switch II that gates phosphate release is widened. This “open door” allows phosphate to exit expeditiously. The converter domain, at the opposite end of the relay helix, is also altered by the increased relay helix flexibility. Simulations show that the region containing the relay helix, activation loop and helix loop helix motif, in the E525K construct shifts towards a more rigor-like conformation that could explain the observed power stroke acceleration. Positioned adjacent to the activation loop, the helix loop helix motif is proposed to form stable interactions with actin during the early stages of actin binding (48). In addition, disruption of the salt bridge interaction between the relay helix and activation loop may allow the loop to interact more easily with actin, further facilitating the actin activation mechanism in the mutant. Steady-state ATPase activity is increased 2-fold in the E525K mutant in the absence of actin. Increased E525K ATPase activity in the absence of actin may result from enhanced relay helix flexibility. Overall, in the absence of the critical salt bridge, actin binding may cause the relay helix to more rapidly transition into intermediate conformations that promote the open back door and post-power stroke lever arm.

### Impact of E525K on IHM formation

A detailed mechanism for how the autoinhibited IHM prevents interaction with actin and thus force generation is currently unclear. Generally, it is understood that the heads of the motor domain fold back asymmetrically onto the S2 rod inhibiting interactions with actin. Our previous work relied on a tarantula myosin homology model (PDB 5TBY) to predict the role of E525K on IHM structural interactions (10, 49). Recently, an updated 3.6 Å cryo-EM model of the human beta-cardiac myosin IHM (PDB 8ACT) has become available (50). The region containing the E525K mutation, the activation loop, and the portion of the lower 50 kDa domain proposed to be involved in the conserved actin activation mechanism are sequestered in the IHM in both models. A model of the IHM based on the 8ACT structure reveals that D900 in the S2 region is in close proximity to E525 in the blocked head (BH) (**Fig. S7**). Thus, the E525K mutation may stabilize the IHM by forming a salt bridge with D900. Overall, we propose that a principal feature of the autoinhibition mechanism of the IHM conformation is that it prevents this region of the myosin motor domain from interacting with actin, subsequently hindering the actin activation mechanism.

### Implications for understanding underlying mechanisms of DCM

Dilated cardiomyopathy *MYH7* mutations are thought to decrease intrinsic motor properties and/or stabilize the autoinhibited state leading to reduced force and power (10, 27, 49, 51). In contrast, HCM mutations are thought to increase intrinsic motor properties and/or destabilize the autoinhibited state leading to increased force and power (23, 52, 53). Interestingly, exceptions are being identified, unraveling the idea that changes in one property of myosin equate to the resulting changes in ensemble force seen in the disease phenotype (10, 19, 20). The E525K DCM mutant is an example of an intriguing exception to this theory showing a DCM mutant can stabilize the autoinhibited state while also increasing intrinsic motor properties along with ensemble force and power of active heads. Ensemble force (*F*_*ensemble*_) can be represented as the product of the number of available heads (*N*) times the duty ratio (*D*).

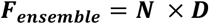

Because E525K mutants present as DCM (↓*F*_*ensemble*_), it is attractive to hypothesize that the ability of E525K to stabilize the autoinhibited state (↓*N*) dominates the increases observed in intrinsic motor properties (↑*D*).

In addition, DCM is known to present with reduced Frank-Starling response, where cardiac muscle has enhanced force generation upon stretch, which makes the tissue less efficient at keeping up with systemic demand (54-59). When muscle is stretched in the presence of load, myosin heads are proposed to be released from the SRX/IHM states (60).This shift allows heads to move away from the thick filament backbone and become readily available to form cross bridges (60). E525K could impair Frank-Starling mediated force increases due to stabilization of the SRX/IHM states, preventing stretch-activation of the thick filament. Because E525K may primarily cause DCM by stabilizing the autoinhibited state, the first inclination would be to treat with an IHM destabilizing therapeutic. However, it is important to note this may trigger the enhanced actin activation mechanism that is concealed by the IHM conformation. Mobilization of the previously sequestered but more powerful motors may lead patients to display hypercontractility and relaxation defects. Further study of the intricately balanced myosin force generating system will continue to elucidate DCM pathogenesis and drive therapeutic development.

### Limitations and Future Directions

It is important to acknowledge any limitations and possible alternative interpretations in the current study. The WT and E525K M2β S1 constructs were examined with endogenous mouse light chains intrinsic to the C2C12 myocyte expression system. This could potentially alter the impact of the E525K mutation, though the location of the mutation makes this unlikely in an S1 construct. In the current study, we were not able to assess the force generating properties of E525K in a simulated thick filament environment. Future studies will perform mechanical measurements with synthetic thick filaments generated with a DNA origami platform (61). Also, utilizing a human induced pluripotent stem cell cardiomyocyte (hiPSC-CM) model would allow for examining the influence of regulatory proteins such as myosin binding protein C.

## Conclusions

In summary, this study revealed an important allosteric pathway associated with actin activation of the myosin power stroke and product release, which we propose is crucial for the actomyosin ATPase mechanism. We suspect that this is a highly conserved allosteric pathway since the critical salt-bridge we identified is conserved throughout most of the myosin superfamily. Our detailed characterization of the E525K mutation in the activation loop also suggests sequestering this element of the actin binding region in the folded-back autoinhibited IHM is crucial for preventing actin activation of the ATP hydrolysis cycle and force generation. Our work further demonstrates the complexity of cardiomyopathy associated mutations in beta-cardiac myosin, since E525K significantly increases duty ratio, ensemble force and power in monomeric subfragment 1 while stabilizing the SRX state and IHM conformation. Thus, there is a delicate balance between intrinsic motor properties of myosin heads recruited to bind the actin thin filament and the ability to autoinhibit myosin heads in the thick filament, which ultimately controls force generation in cardiac muscle. The severity of the cardiomyopathy phenotype, whether it be the enhanced contractile force and disrupted relaxation observed in HCM or the depressed force observed in DCM, will depend on the interplay between these two key factors. Thus, detailed biophysical characterization of cardiomyopathy associated mutations in sarcomeric proteins will help guide future therapies that attempt to normalize these key factors that are crucial for controlling cardiac contractility.

## Materials and Methods

Detailed materials and methods are outlined in *SI Appendix, Materials and Methods*.

## Supporting information

Bodt et al. 2023

## Acknowledgments

C.M.Y. is supported by NIH grants R01 HL163585 and R01HL150953. W.M. and J.A.M. were supported by NIH grant R01GM031749. The authors gladly acknowledge the computational resources (Triton Shared Computing Cluster, 10.57873/T34W2R) provided by San Diego Supercomputer Center at the University of California, San Diego.

## Supporting Information

### Materials and Methods

#### Reagents

MOPS 5/4 buffer was used for all kinetic solution experiments (10 mM MOPS, 5 mM KCL, 1 mM EGTA, 4 mM MgCl_2_, and 1 mM dithiothreitol [DTT], pH 7.0). Both ATP and ADP were prepared from lyophilized powder and concentrations of each were determined by absorbance at 259 nm (ε259 = 15,400 M^−1^·cm^−1^). Cy3ATP/ADP was purchased from Jena Biosciences. 2′-deoxy-ATP/ADP labeled with N-methylanthraniloyl at the 3′-ribose position (mant*-*ADP) were also purchased from Jena Biosciences. Alexa Flour 488 Maleimide powder was purchased from Invitrogen and solubilized in dimethyl sulfoxide (DMSO) for RLC labeling. Fluorescently labeled phosphate binding protein (MDCC-PBP) was prepared as described (1).

#### Protein construction, expression, and purification

The M2β S1 cDNA sequence from the *MYH7* gene (GenBank: AAA51837.1) containing a C-terminal Avi tag (GLNDIFEAQKIEWHE) and FLAG (DYKDDDDK) tag was cloned into the pDual shuttle vector. A M2β S1 GFP construct containing a C-terminal GFP tag followed by Avi tag and FLAG tag was also generated for motility assays. The E525K mutation was engineered into these constructs using the Quikchange site-directed mutagenesis (Agilent). Recombinant adenovirus was prepared by Vector Biolabs (Malvern, Pa) at a titer of 10^8^ plaque forming units per ml (pfu/ml). The virus was expanded by infecting Ad293 cells at a multiplicity of infection (MOI) of 3-5 (2). The virus was harvested and purified by CsCl density sedimentation, giving a final virus titer of 10^10^-10^11^. Mouse skeletal muscle derived C2C12 cells were used to express the cardiac myosin constructs as previously described (2-5). Briefly, cells were grown to ∼90% confluency on (145/20mm) tissue culture plates in growth media. Media on 20 plates was changed to fusion media and virus at 4 × 10^7^ pfu/ml on the day of infection. Protein was purified with FLAG affinity chromatography from cells harvested 10-12 days post-infection. For power stroke FRET, endogenous mouse regulatory light chain (RLC) was replaced with Alexa 488-labeled human RLC (A488RLC) on M2β S1 using an on-column exchange method described previously (6, 7). Actin was purified from rabbit skeletal muscle acetone power (Pellfreeze) (8). Actin was labeled with pyrene iodoacetamide for transient kinetic experiments when required.

#### Steady-state ATPase measurements

The steady-state actin-activated ATPase activity of 0.1 μM M2β S1 was measured at varying actin concentrations (0, 5, 10, 20, 40, and 60 μM) utilizing an NADH-coupled ATP regenerating system. Steady-state ATPase was monitored for 200 s at 25°C at 0.2 s intervals which was then plotted as a function of actin concentration and fitted to the Michaelis-Menten equation to calculate the *k*_cat_ and *K*_ATPase_ (9, 10)

#### Single turnover measurements

Fluorescence of mant-ATP was monitored with 290 nm excitation and 395 nm long-pass emission filter. Myosin was incubated on ice for 10 minutes in appropriate buffer conditions before each experiment. Single mant-ATP turnover experiments were performed by first incubating M2β S1 with mant-ATP (1 μM) for 30 s at room temperature followed by mixing the complex with saturating ATP (2 mM). Fluorescence decay was monitored for 1000 s and SRX ratio, SRX rate, and DRX rate were determined (11).

#### Transient kinetic measurements

All transient kinetic experiments were performed with the Applied Photophysics stopped-flow device equipped with an excitation monochromator, 1.2 ms dead-time, and a 9.3 nm band pass. Tryptophan fluorescence for ATP binding and hydrolysis experiments was excited at 290 nm and monitored with a 320 nm long-pass emission filter. Experiments with pyrene actin fluorescence were monitored with 365 nm excitation and a 395 nm long-pass emission filter. Mant fluorescence was monitored with 290 nm excitation and a 395 nm long-pass emission filter. MDCC-PBP in phosphate release experiments was monitored with a 380 nm excitation and a 425 nm long-pass emission filter. The stopped-flow software program was utilized to fit fluorescent transients.

#### Power stroke FRET measurements

Change in donor fluorescence (A488RLC) using an excitation wavelength of 460 nm and an interference filter (500-525 nm) to eliminate background from Cy3ATP fluorescence was examined by stopped flow (6).

#### *In Vitro* motility assays

The *in vitro* motility assay was performed with WT and E525K M2β S1 constructs containing a C-terminal GFP tag, in accordance with established protocols (11, 12). Briefly, coverslips were coated with 0.2% nitrocellulose in amyl acetate (Ladd Research) and adhered to microscope slides via double-sided tape creating a flow cell. For density dependence measurements, GFP antibody (0.1 mg/ml, Thermo Fisher G10362) was directly applied to the nitrocellulose surface and subsequently blocked by BSA (1 mg/ml) before the addition of myosin at concentrations (0.05-0.7 μM) in MOPS 50 buffer (10 mM MOPS, pH 7.0, 50 mM KCl, 1 mM EGTA, 1 mM MgCl_2_, 1 mM DTT). To ensure inactive myosin heads on the surface were blocked, unlabeled sheared actin (2 μM) was added and chased with ATP (2 mM). Actin labeled with Alexa 555-phalloidin (10 nM) was then added followed by an activation buffer containing methylcellulose (0.35%), ATP regeneration system (2 mM ATP, 5 mg mL^−1^ glucose, 45 units mL^−1^ pyruvate kinase, and 0.45 mM phosphoenolpyruvate), and oxygen scavengers (0.1 mg mL^−1^ glucose oxidase and 0.018 mg mL^−1^ catalase) to initiate motility. The frictional loading assay was further executed by mixing GFP antibody with alpha-actinin (Cytoskeleton AT01) at concentrations ranging from 0-22.5 ug/mL until actin filaments exhibited cessation of movement. Subsequently, myosin was introduced into the MOPS 50 buffer at a constant concentration of 0.4 µM for all frictional test samples. Motility was observed utilizing a Leica DMi8 fluorescence microscope equipped with a 63×/1.4 oil objective and DsRed filter, with excitation/emission 555/620 nm. Image acquisition was performed at 1 s intervals for a total duration of 120 s. The videos were exported to ImageJ for manual analysis, compiling with three different protein preparations to facilitate statistical comparing between WT and E525K.

#### Model and parameters for frictional loaded assay

The surface concentration of alpha-actinin, in units of molecules per area, is given by:

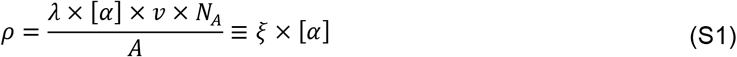

where *λ* is the fraction of alpha-actinin that bound to the nitrocellulose surface, *λ* = 6% was approximately measured under similar conditions (13). *ν* = 10 *µL* is the volume of alpha-actinin added, [*α*] is the molar concentration of alpha-actinin added to the flow cell, *N*_*A*_ is Avogadro’s number and *A* is the surface area of the flow cell, dimension *Width* × *Length* = 5 *mm* × 22 *mm, A* = 110 *mm*^2^ = 1.1 × 10^−4^ *m*^2^.

The frictional load imposed by the bed of alpha-actinin molecules would be given by:

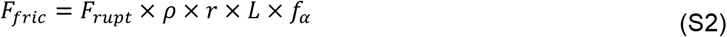

*F*_*rupt*_ is the rupture force, *r* is the reach of alpha-actinin to bind to actin filament. Considering the length of an alpha-actinin molecule 40 nm (14, 15) and an actin filament in a motility assay is 10 nm above the flow cell surface, the distance over which an alpha-actinin can reach and attach to actin *r* = 82 *nm. L* is the average length of an actin filament. It is noteworthy that the presence of alpha-actinin can lead to variations in actin filament length, potentially inducing a shredding effect. Average actin filament length was quantified based on an average of 20 filaments per condition.

*f*_*α*_ is the fraction of alpha-actinin lifetime spent attached to actin.

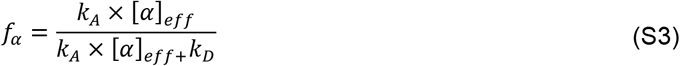

*k*_*A*_ = 4 × 10^6^ *M*^−1^*s*^−1^ is the second-order rate constant for alpha-actinin attachment to actin, *k*_*D*_ = 9.6 *s*^−1^ is the alpha-actinin detachment rate, [*α*]_*eff*_ is the effective molarity of alpha-actinin on the surface, 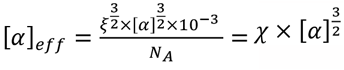.

The velocity of actin filament sliding *V* as a function alpha-actinin concentration [*α*] and fitted to an elastic model (16) shown in **Fig. 5B**.

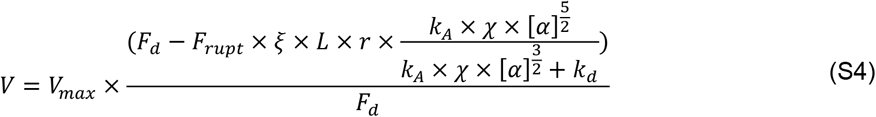

with the fitting parameters of myosin driving force, *F*_*d*_, and maximal unloaded sliding velocity, *V*_*max*_.

Using Eq.S2, alpha-actinin concentrations were converted to forces, and the data were fit to the Hill equation (**Fig. 5C**).

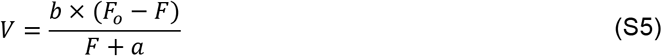

Power was then converted by *P* = *F* × *V*, and fit to the Hill equation (**Fig. 5D**).

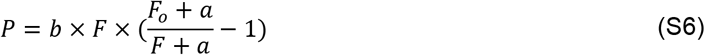

#### Kinetic modeling

A kinetic model that included all major forward and reverse rate constants was generated (see **Table S1**) and used to produce simulations of steady-state ATPase activity using Kintek Explorer. Fluorescence transients of the power stroke and phosphate release were also fit to the proposed kinetic model.

#### Gaussian accelerated molecular dynamics simulations

The initial structures of the E525K actomyosin complex were obtained from a previous work (17). For each Rosetta model of the PPS actomyosin, E525 was mutated into a lysine residue and three independent replica GaMD runs were performed as described (17). Each E525K model was solvated in a water box with 150 mM NaCl. All the simulations were performed using the GPU version of Amber18 with the ff14SB force field (18, 19). Firstly, energy minimization was carried out with harmonic restraints applied to protein atoms using a spring constant of 1 kcal/(mol Å^2^). Keeping the same positional constraints, a subsequent 5 ns equilibration MD was performed at 300 K, while the temperature was maintained by Langevin dynamics with a friction coefficient of 1 ps^−1^. Next, Gaussian accelerated MD (GaMD) was employed to enhance the sampling of protein conformational dynamics (20). A 5-ns conventional MD stage was first used to gather statistics for determining the initial GaMD acceleration parameters. Then a 20-ns GaMD stage was conducted under isothermal-isobaric conditions at 1 bar and 300 K. Both total potential energy boost and dihedral energy boost were applied to the system, each having a 6 kcal/mol upper limit of the standard deviation to ensure accurate reweighting. The accumulated GaMD simulation time (72 trajectories in total) amounted to 1.44 µs. The free energy profile was obtained using the reweighting approach detailed in *Ma et al*. (17). The analyses for the WT case were based on our previous GaMD simulations of the WT PPS actomyosin (17).

#### Statistics

One biological replicate is defined as an independent protein preparation and technical replicates are defined as repeated measurements from the same protein preparation. Direct comparisons were made between WT and E525K protein prepared in parallel. Sample sizes refer to biological replicates with a minimum of (N=3) for all experiments unless otherwise noted. Unpaired Student’s t-tests were performed to compare differences between groups in all experiments with statistical error reported.

**Table S1.**
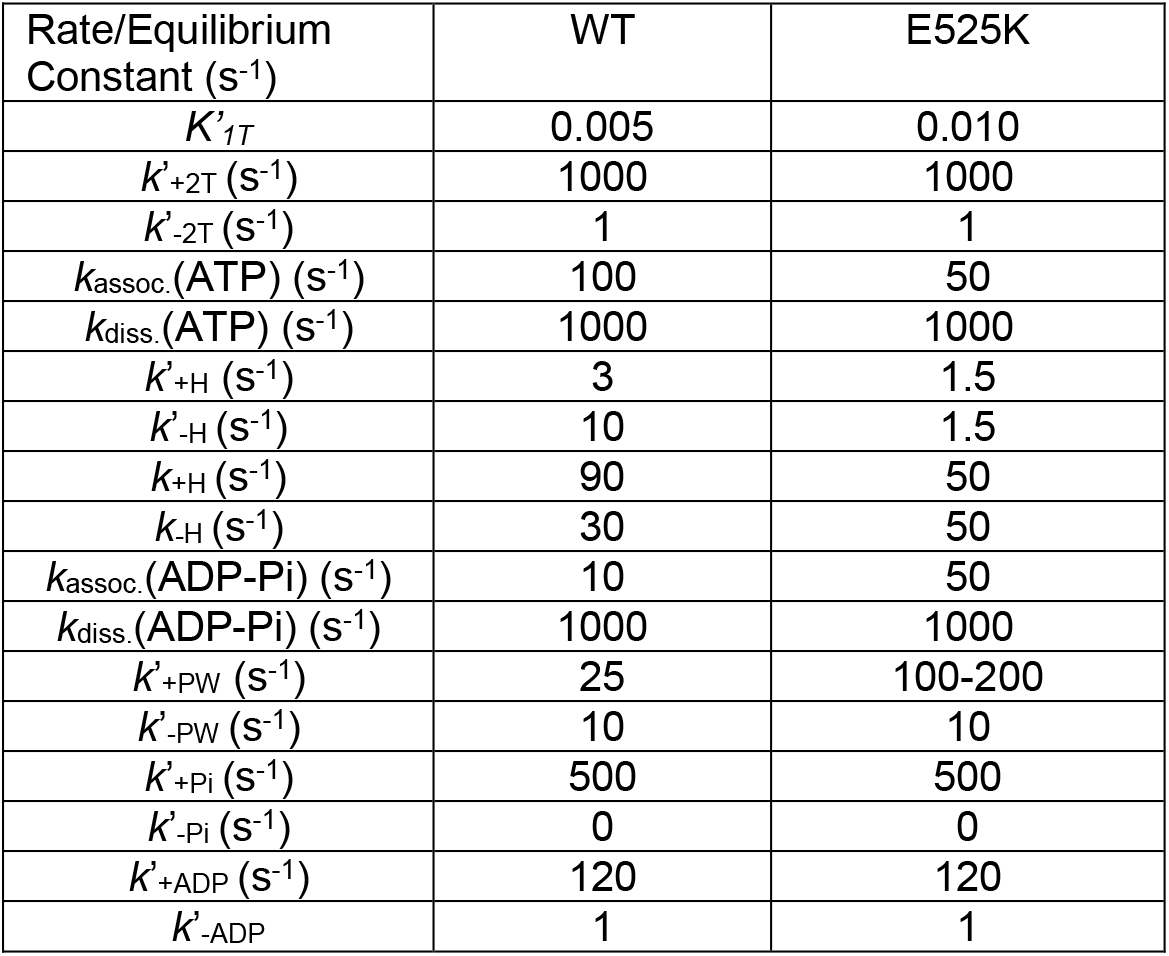
Rate and equilibrium constants for simulations. Rate and equilibrium constants used to simulate the steady-state ATPase as well as the power stroke and phosphate release rate constants (**Fig. S5**).

| Rate/Equilibrium Constant (s<sup>-1</sup>) | WT | E525K |
| --- | --- | --- |
| $K'_{1T}$ | 0.005 | 0.010 |
| $k'_{+2T}$ (s<sup>-1</sup>) | 1000 | 1000 |
| $k'_{-2T}$ (s<sup>-1</sup>) | 1 | 1 |
| $k_{\text{assoc.}}(\text{ATP})$ (s<sup>-1</sup>) | 100 | 50 |
| $k_{\text{diss.}}(\text{ATP})$ (s<sup>-1</sup>) | 1000 | 1000 |
| $k'_{+H}$ (s<sup>-1</sup>) | 3 | 1.5 |
| $k'_{-H}$ (s<sup>-1</sup>) | 10 | 1.5 |
| $k_{+H}$ (s<sup>-1</sup>) | 90 | 50 |
| $k_{-H}$ (s<sup>-1</sup>) | 30 | 50 |
| $k_{\text{assoc.}}(\text{ADP-Pi})$ (s<sup>-1</sup>) | 10 | 50 |
| $k_{\text{diss.}}(\text{ADP-Pi})$ (s<sup>-1</sup>) | 1000 | 1000 |
| $k'_{+PW}$ (s<sup>-1</sup>) | 25 | 100-200 |
| $k'_{-PW}$ (s<sup>-1</sup>) | 10 | 10 |
| $k'_{+Pi}$ (s<sup>-1</sup>) | 500 | 500 |
| $k'_{-Pi}$ (s<sup>-1</sup>) | 0 | 0 |
| $k'_{+ADP}$ (s<sup>-1</sup>) | 120 | 120 |
| $k'_{-ADP}$ | 1 | 1 |

**Figure S1.**
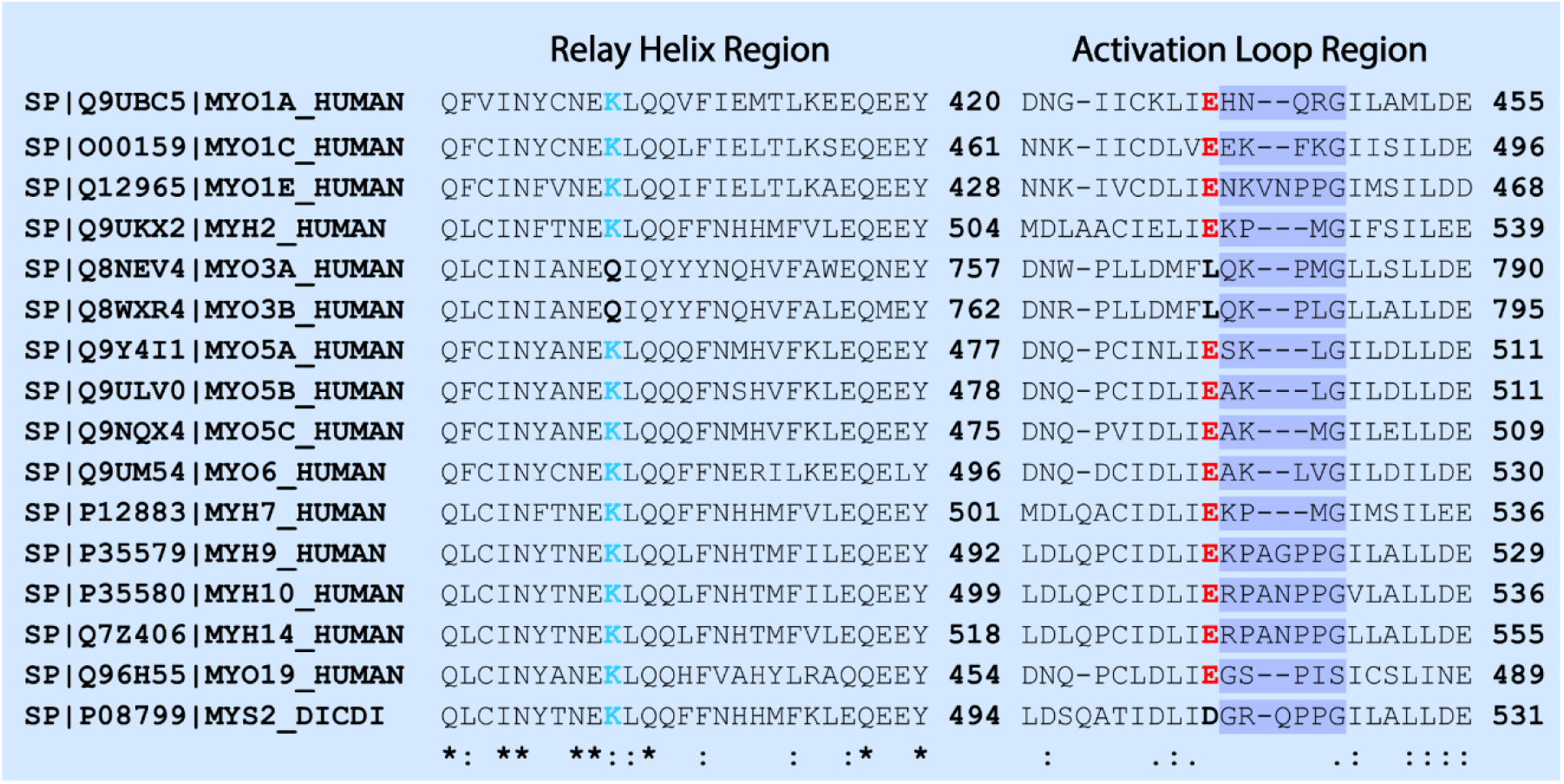
Alignment of relay helix and activation loop regions. *Uniprot* sequences were aligned using the *Clustal Omega* multiple sequence alignment program. An * (asterisk) indicates positions which have a single, fully conserved residue. A : (colon) indicates conservation between groups of strongly similar properties - roughly equivalent to scoring > 0.5 in the Gonnet PAM 250 matrix: STA. A. (period) indicates conservation between groups of weakly similar properties - roughly equivalent to scoring =< 0.5 and > 0 in the Gonnet PAM 250 matrix. The activation loop region was highlighted (dark blue) based on the length of the MYH7 loop. Lysine (light blue) in the relay helix region (K484 in cardiac) and glutamate (red) in the activation loop region (E525 in cardiac) are highly conserved across myosin isoforms. These residues are essential for the proposed allosteric actin activation mechanism. MYO3A and MYO3B were identified as exceptions with glutamine and leucine residues in place of lysine and glutamate respectively.

**Figure S2.**
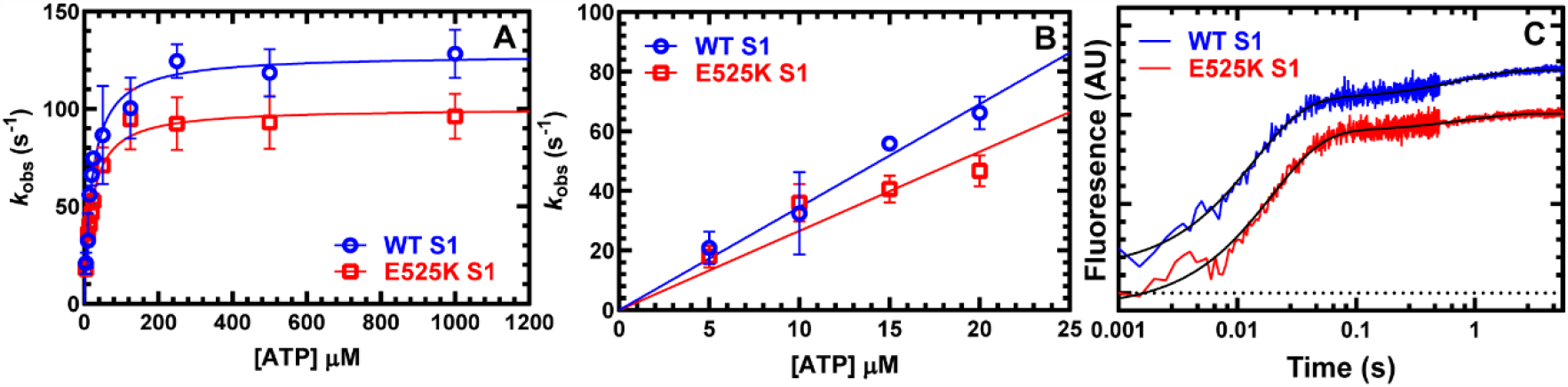
ATP binding and hydrolysis in myosin. Tryptophan fluorescence enhancement was used to monitor ATP binding/hydrolysis by mixing 1 μM M2β S1 WT or E525K with varying concentrations of ATP. The fluorescence transients were fit to a double exponential function. **(A)** The observed fast phase was plotted as a function of ATP concentration and fit to a hyperbolic function to determine maximum rate of ATP hydrolysis. **(B)** The second-order rate constant for ATP binding was determined by the linear fit at low ATP concentrations. **(C)** Representative fluorescence transients (average of 2 transients) in the presence of 25 μM ATP are shown fit to a double exponential function. Data points at each ATP concentration represent the average ± SD of three experiments from three separate protein preparations.

**Figure S3.**
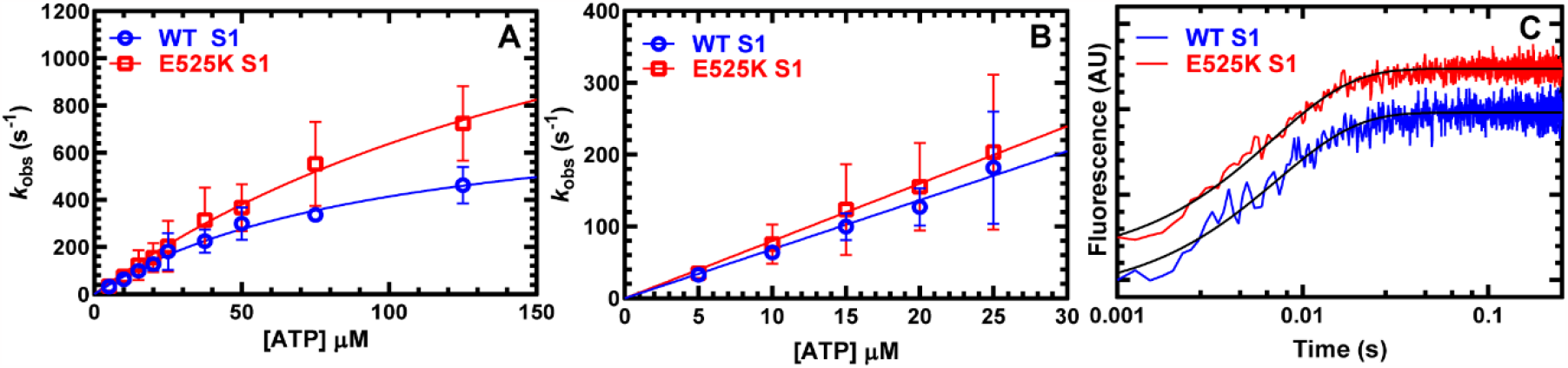
ATP binding to actomyosin. ATP-induced dissociation from pyrene actin was performed by mixing a complex of M2β S1:pyrene actin (0.5 μM M2β S1 and pyrene actin) with a series of ATP concentrations (5 to 250 μM). The fluorescence transients were fit to a single exponential function at all ATP concentrations. **(A)** The observed rate constant was hyperbolically dependent on ATP concentrations. **(B)** The second-order rate constant for ATP binding was determined from the linear fit at low ATP concentrations. **(C)** Representative fluorescence transients (average of 2 transients) in the presence of 25 μM ATP are shown fit to a single exponential function. Data points at each actin concentration represent the average ± SD of five experiments from five separate protein preparations.

**Figure S4.**
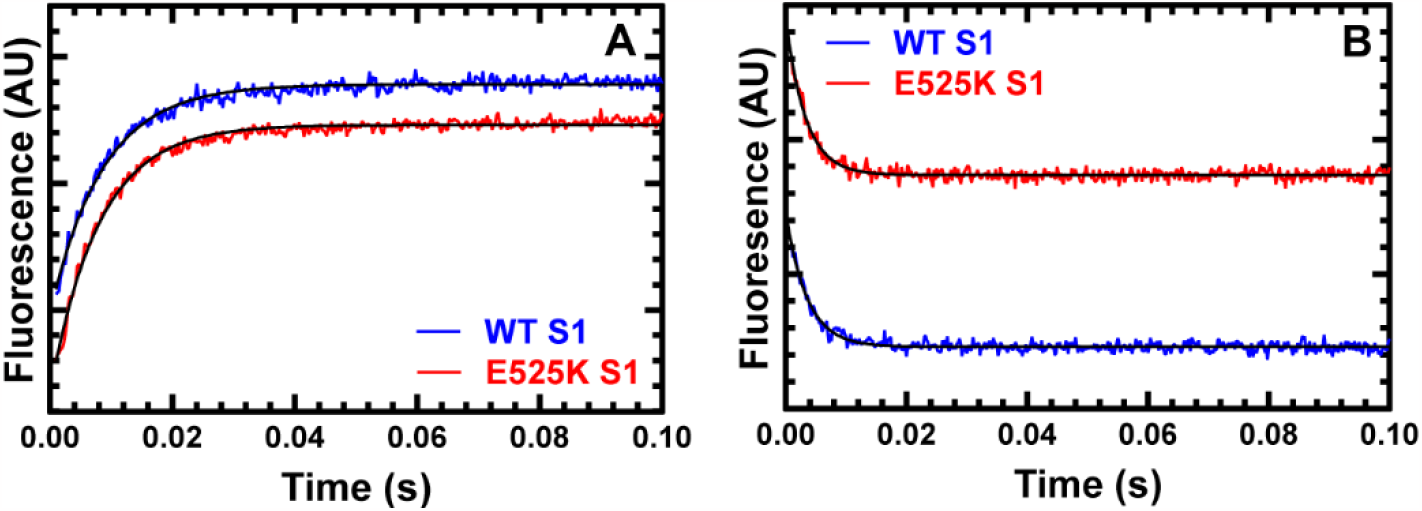
ADP release from acto-myosin. **(A)** The ADP release rate constant was examined by mixing pyrene actomyosin.ADP (0.5 μM M2β S1 and pyrene actin, 10 μM ADP) with saturating ATP (2 mM). Fluorescent transients were fit to a single exponential function. **(B)** The ADP release rate constant was also examined with mant-labeled ADP by mixing a complex of 0.5 μM M2β S1 and actin, and 10 μM mantADP with 1 mM ATP. Representative fluorescence transients of mant-ADP release from actomyosin (average of 2 transients) fit to a single exponential function are shown.

**Figure S5.**
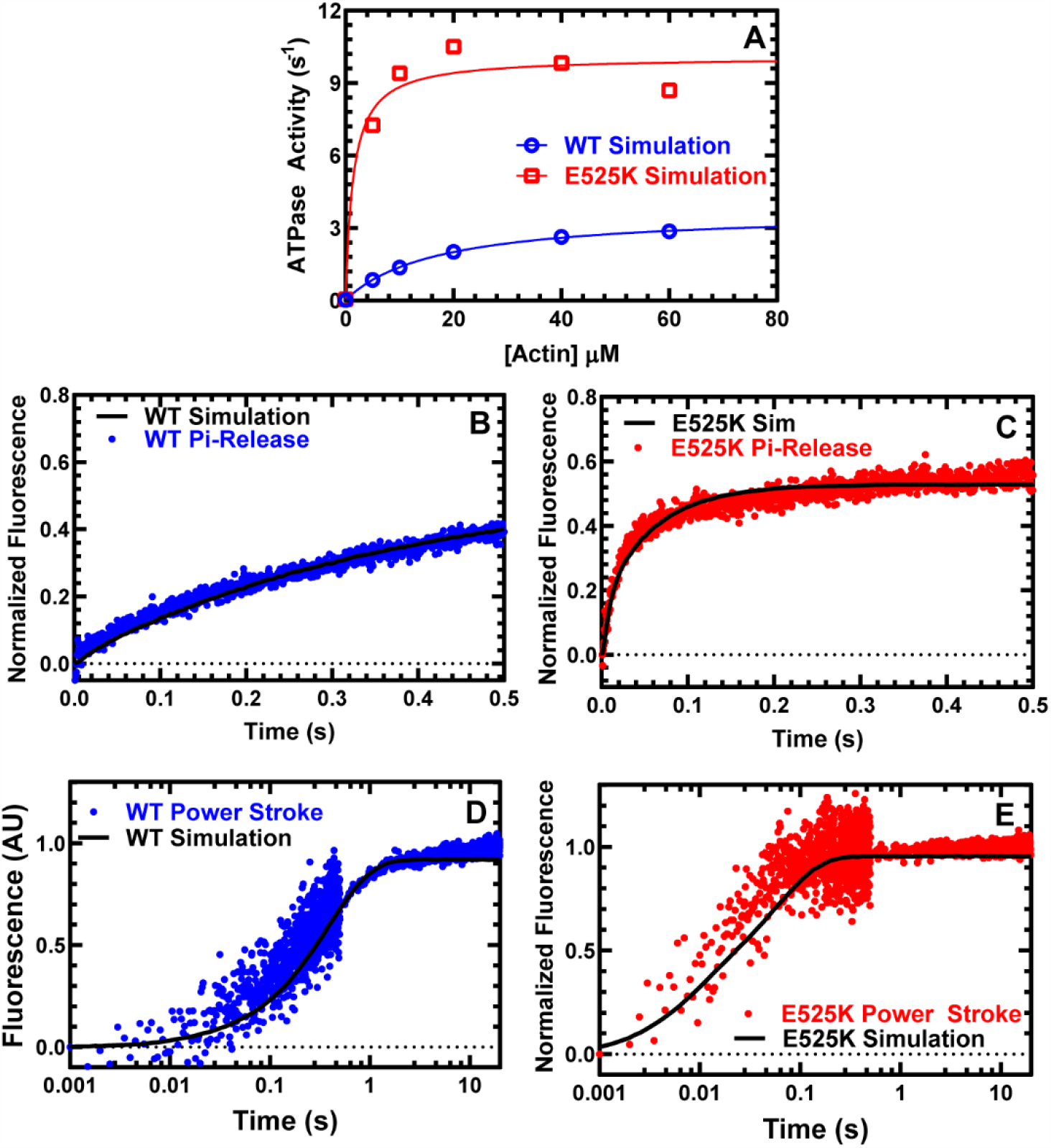
Kinetic simulations. **(A)** The rate and equilibrium constants determined experimentally and estimated from previous studies (**see Table S1**) were used to simulate the steady-state ATPase for both WT and E525K. **(B & C)** The phosphate release transients and **(D & E)** power stroke transients were compared to simulated data in WT and E525K, respectively. Only the first 0.5 sec of the Pi-release transients were simulated to focus on the initial Pi-burst associated with actin-activated Pi-release.

**Figure S6.**
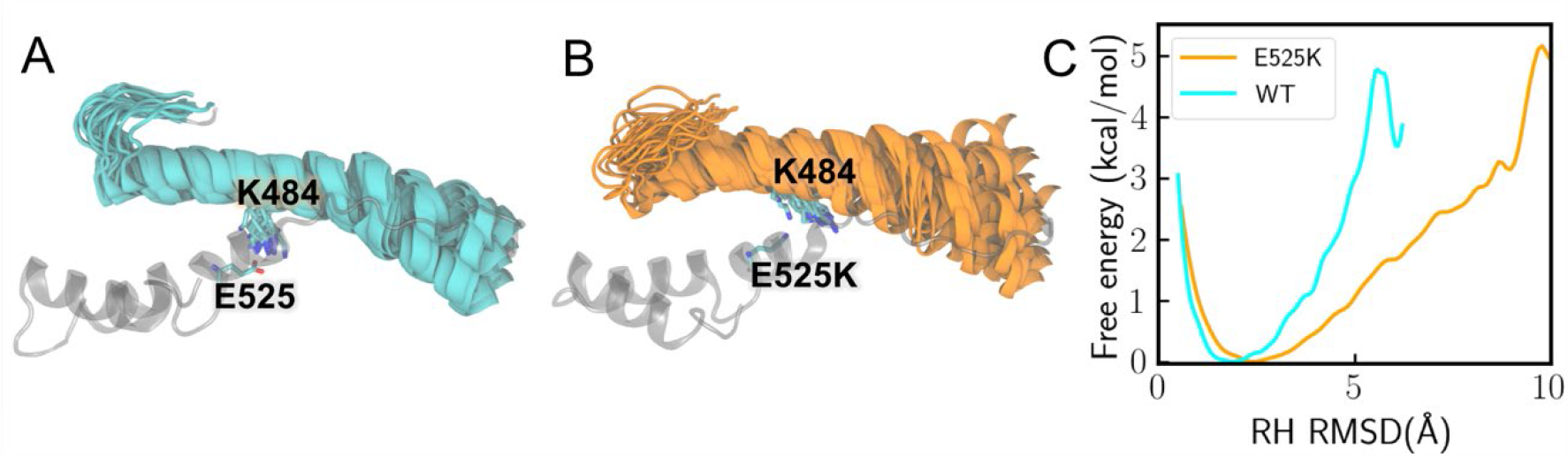
Enhanced flexibility of the relay helix in the E525K simulations. **(A) (B)** All the MD clusters of the relay helix (RH) are overlaid after being aligned to residues 519 to 540 (WT in cyan; E525K in orange). Variations in the K484 side chain are also demonstrated in both panels. **(C)** The free energy profile is plotted along the RH RMSD coordinate. E525K is much more likely than WT to exhibit RH RMSD values greater than 3Å.

**Figure S7.**
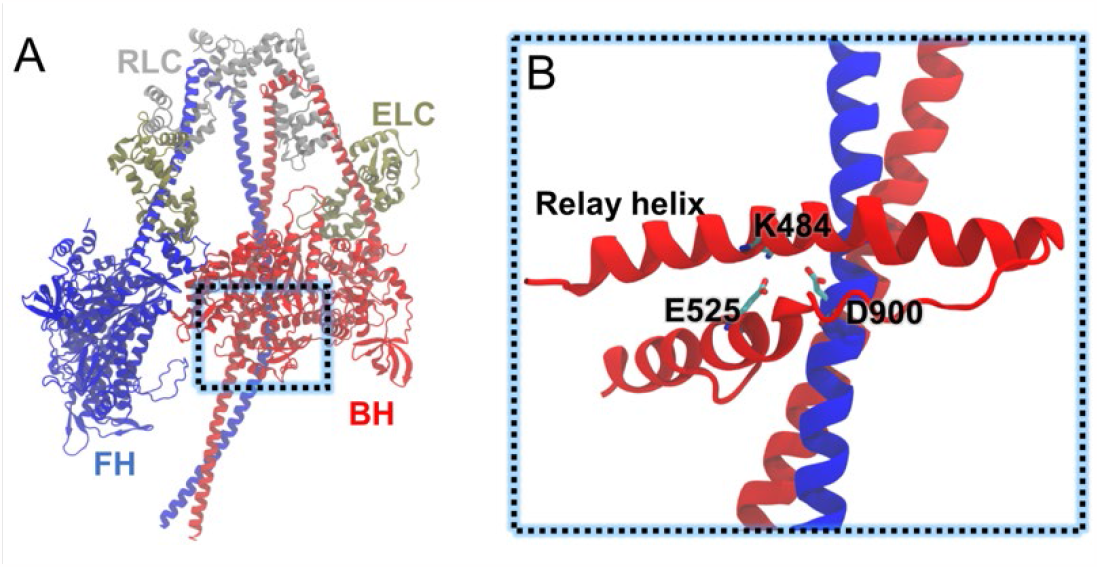
Model to explain how E525K may stabilize IHM formation. By forming an additional salt bridge (E525K and D900), the mutation E525K could stabilize the interface between the blocked head (BH-red) and subfragment-2 of the free head (FH-blue). The 8ACT PDB structure was used to generate the IHM model.

## Notes

### Competing Interest Statement

The authors have declared no competing interest.

