## Supplementary material for "Dilated cardiomyopathy mutation in beta-cardiac myosin enhances actin activation of the power stroke and phosphate release": Bodt et al. 2023

Christopher M. Yengo

**This PDF file includes:**

Materials and Methods
Table S1. Rate and equilibrium constants for simulations.
Figure S1. Alignment of relay helix and activation loop regions.
Figure S2. ATP binding and hydrolysis in myosin.
Figure S3. ATP binding to actomyosin.
Figure S4. ADP release from acto-myosin.
Figure S5. Kinetic simulations.
Figure S6. Enhanced flexibility of the relay helix in the E525K simulations.
Figure S7. Model to explain how E525K may stabilize IHM formation.
Supporting Information References

### Model and parameters for frictional loaded assay

The surface concentration of alpha-actinin, in units of molecules per area, is given by:

$$\rho = \frac{\lambda \times [\alpha] \times v \times N_A}{A} \equiv \xi \times [\alpha] \qquad \text{(S1)}$$

where $\lambda$ is the fraction of alpha-actinin that bound to the nitrocellulose surface, $\lambda = 6\%$ was approximately measured under similar conditions (13). $v = 10\ \mu L$ is the volume of alpha-actinin added,
$[\alpha]$ is the molar concentration of alpha-actinin added to the flow cell, $N_A$ is Avogadro's number and $A$ is the surface area of the flow cell, dimension $Width \times Length = 5\ mm \times 22\ mm$, $A = 110\ mm^2 = 1.1 \times 10^{-4}\ m^2$ .
The frictional load imposed by the bed of alpha-actinin molecules would be given by:

$$F_{fric} = F_{rupt} \times \rho \times r \times L \times f_\alpha \qquad \text{(S2)}$$

$F_{rupt}$ is the rupture force, $r$ is the reach of alpha-actinin to bind to actin filament. Considering the length of an alpha-actinin molecule 40 nm (14, 15) and an actin filament in a motility assay is 10 nm above the flow cell surface, the distance over which an alpha-actinin can reach and attach to actin $r = 82\ nm.$ $L$ is the average length of an actin filament. It is noteworthy that the presence of alpha-actinin can lead to variations in actin filament length, potentially inducing a shredding effect. Average actin filament length was quantified based on an average of 20 filaments per condition.
$f_\alpha$ is the fraction of alpha-actinin lifetime spent attached to actin.

$$f_\alpha = \frac{k_A \times [\alpha]_{eff}}{k_A \times [\alpha]_{eff+} k_D} \qquad \text{(S3)}$$

$k_A = 4 \times 10^6\ M^{-1}s^{-1}$ is the second-order rate constant for alpha-actinin attachment to actin, $k_D = 9.6\ s^{-1}$ is the alpha-actinin detachment rate, $[\alpha]_{eff}$ is the effective molarity of alpha-actinin on the surface, $[\alpha]_{eff} = \frac{\xi^{\frac{3}{2}} \times [\alpha]^{\frac{3}{2}} \times 10^{-3}}{N_A} = \chi \times [\alpha]^{\frac{3}{2}}$.
The velocity of actin filament sliding $V$ as a function alpha-actinin concentration $[\alpha]$ and fitted to an elastic model (16) shown in **Fig. 5B**.

$$V = V_{max} \times \frac{(F_d - F_{rupt} \times \xi \times L \times r \times \frac{k_A \times \chi \times [\alpha]^{\frac{5}{2}}}{k_A \times \chi \times [\alpha]^{\frac{3}{2}} + k_d})}{F_d} \qquad \text{(S4)}$$

with the fitting parameters of myosin driving force, $F_d$ , and maximal unloaded sliding velocity, $V_{max}$.
Using Eq.S2, alpha-actinin concentrations were converted to forces, and the data were fit to the Hill equation (**Fig. 5C**).

$$V = \frac{b \times (F_o - F)}{F + a} \qquad \text{(S5)}$$

Power was then converted by $P = F \times V$, and fit to the Hill equation (**Fig. 5D**).

$$P = b \times F \times (\frac{F_o + a}{F + a} - 1) \qquad \text{(S6)}$$

**Kinetic modeling.**
A kinetic model that included all major forward and reverse rate constants was generated (see **Table S1**) and used to produce simulations of steady-state ATPase activity using Kintek Explorer. Fluorescence transients of the power stroke and phosphate release were also fit to the proposed kinetic model.

**Table S1. Rate and equilibrium constants for simulations**. Rate and equilibrium constants used to simulate the steady-state ATPase as well as the power stroke and phosphate release rate constants (**Fig. S5**).

| Rate/Equilibrium Constant ($s^{-1}$) | WT | E525K |
|---|---|---|
| $K'_{1T}$ | 0.005 | 0.010 |
| $k'_{+2T}$ ($s^{-1}$) | 1000 | 1000 |
| $k'_{-2T}$ ($s^{-1}$) | 1 | 1 |
| $k_{assoc.}$(ATP) ($s^{-1}$) | 100 | 50 |
| $k_{diss.}$(ATP) ($s^{-1}$) | 1000 | 1000 |
| $k'_{+H}$ ($s^{-1}$) | 3 | 1.5 |
| $k'_{-H}$ ($s^{-1}$) | 10 | 1.5 |
| $k_{+H}$ ($s^{-1}$) | 90 | 50 |
| $k_{-H}$ ($s^{-1}$) | 30 | 50 |
| $k_{assoc.}$(ADP-Pi) ($s^{-1}$) | 10 | 50 |
| $k_{diss.}$(ADP-Pi) ($s^{-1}$) | 1000 | 1000 |
| $k'_{+PW}$ ($s^{-1}$) | 25 | 100-200 |
| $k'_{-PW}$ ($s^{-1}$) | 10 | 10 |
| $k'_{+Pi}$ ($s^{-1}$) | 500 | 500 |
| $k'_{-Pi}$ ($s^{-1}$) | 0 | 0 |
| $k'_{+ADP}$ ($s^{-1}$) | 120 | 120 |
| $k'_{-ADP}$ | 1 | 1 |

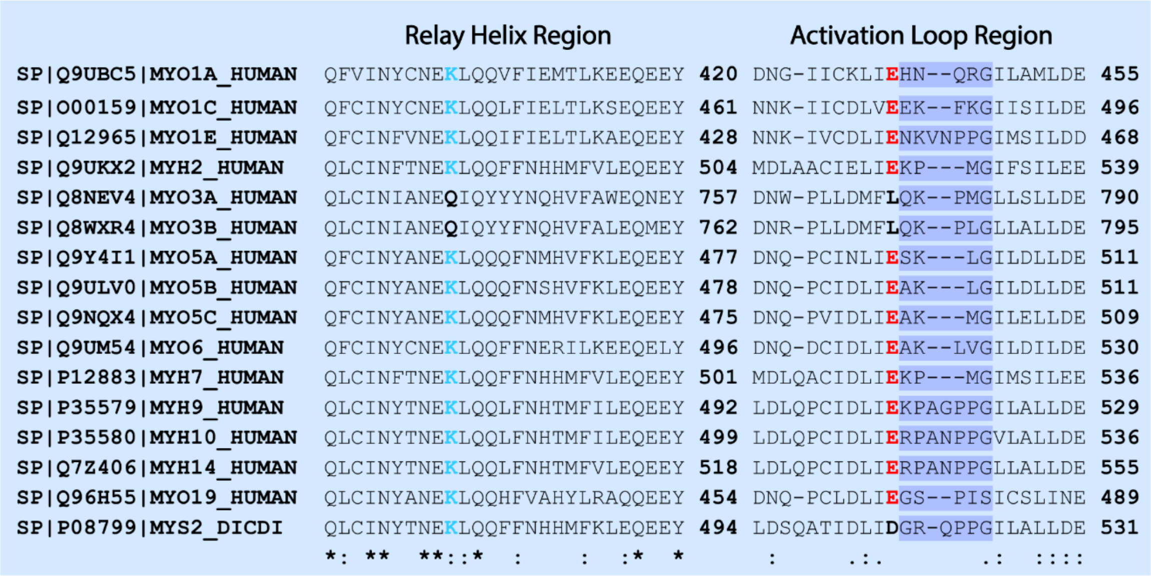


**Figure S1. Alignment of relay helix and activation loop regions.** *Uniprot* sequences were aligned using the *Clustal Omega* multiple sequence alignment program. An * (asterisk) indicates positions which have a single, fully conserved residue. A : (colon) indicates conservation between groups of strongly similar properties - roughly equivalent to scoring > 0.5 in the Gonnet PAM 250 matrix: STA. A . (period) indicates conservation between groups of weakly similar properties - roughly equivalent to scoring =< 0.5 and > 0 in the Gonnet PAM 250 matrix. The activation loop region was highlighted (dark blue) based on the length of the MYH7 loop. Lysine (light blue) in the relay helix region (K484 in cardiac) and glutamate (red) in the activation loop region (E525 in cardiac) are highly conserved across myosin isoforms. These residues are essential for the proposed allosteric actin activation mechanism. MYO3A and MYO3B were identified as exceptions with glutamine and leucine residues in place of lysine and glutamate respectively.

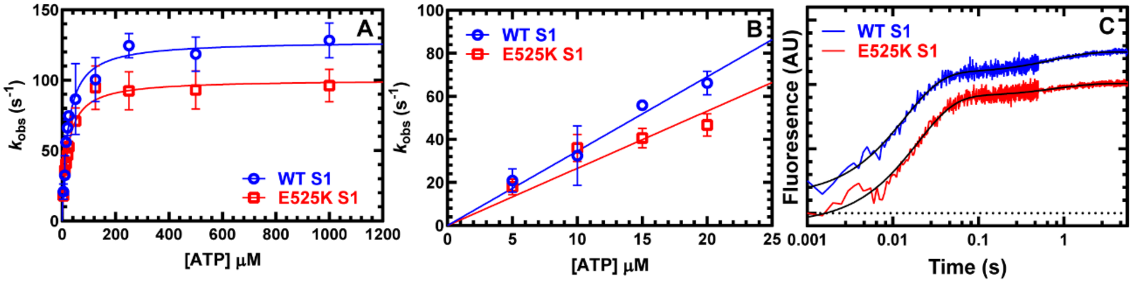


**Figure S2. ATP binding and hydrolysis in myosin.** Tryptophan fluorescence enhancement was used to monitor ATP binding/hydrolysis by mixing 1 μM M2β S1 WT or E525K with varying concentrations of ATP. The fluorescence transients were fit to a double exponential function. **(A)** The observed fast phase was plotted as a function of ATP concentration and fit to a hyperbolic function to determine maximum rate of ATP hydrolysis. **(B)** The second-order rate constant for ATP binding was determined by the linear fit at low ATP concentrations. **(C)** Representative fluorescence transients (average of 2 transients) in the presence of 25 μM ATP are shown fit to a double exponential function. Data points at each ATP concentration represent the average ± SD of three experiments from three separate protein preparations.

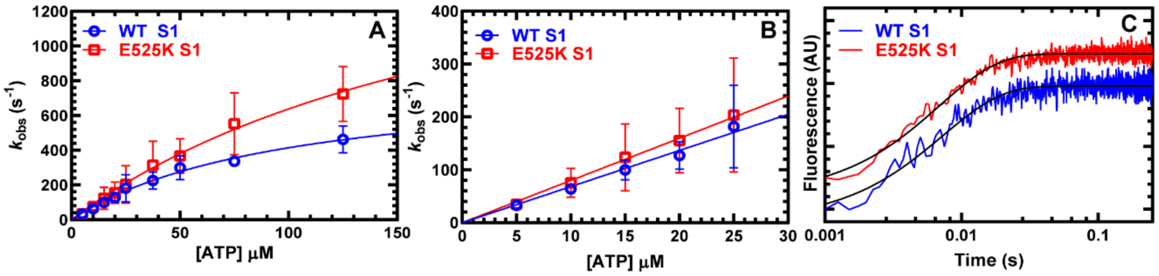


**Figure S3. ATP binding to actomyosin.** ATP-induced dissociation from pyrene actin was performed by mixing a complex of M2β S1:pyrene actin (0.5 µM M2β S1 and pyrene actin) with a series of ATP concentrations (5 to 250 µM). The fluorescence transients were fit to a single exponential function at all ATP concentrations. **(A)** The observed rate constant was hyperbolically dependent on ATP concentrations. **(B)** The second-order rate constant for ATP binding was determined from the linear fit at low ATP concentrations. **(C)** Representative fluorescence transients (average of 2 transients) in the presence of 25 µM ATP are shown fit to a single exponential function. Data points at each actin concentration represent the average ± SD of five experiments from five separate protein preparations.

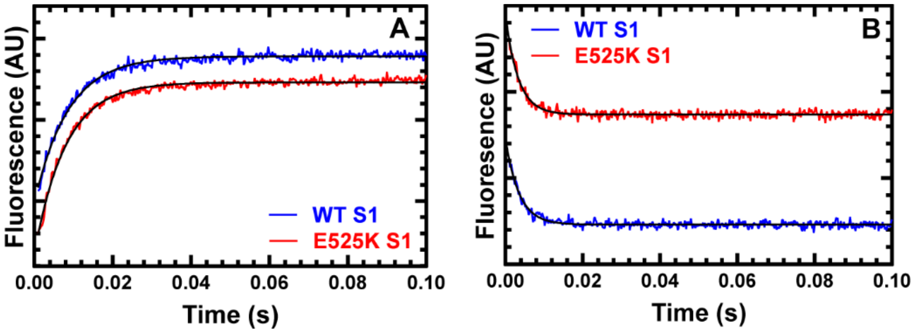


**Figure S4. ADP release from acto-myosin. (A)** The ADP release rate constant was examined by mixing pyrene actomyosin.ADP (0.5 μM M2β S1 and pyrene actin, 10 μM ADP) with saturating ATP (2 mM). Fluorescent transients were fit to a single exponential function. **(B)** The ADP release rate constant was also examined with mant-labeled ADP by mixing a complex of 0.5 μM M2β S1 and actin, and 10 μM mantADP with 1 mM ATP. Representative fluorescence transients of mant-ADP release from actomyosin (average of 2 transients) fit to a single exponential function are shown.

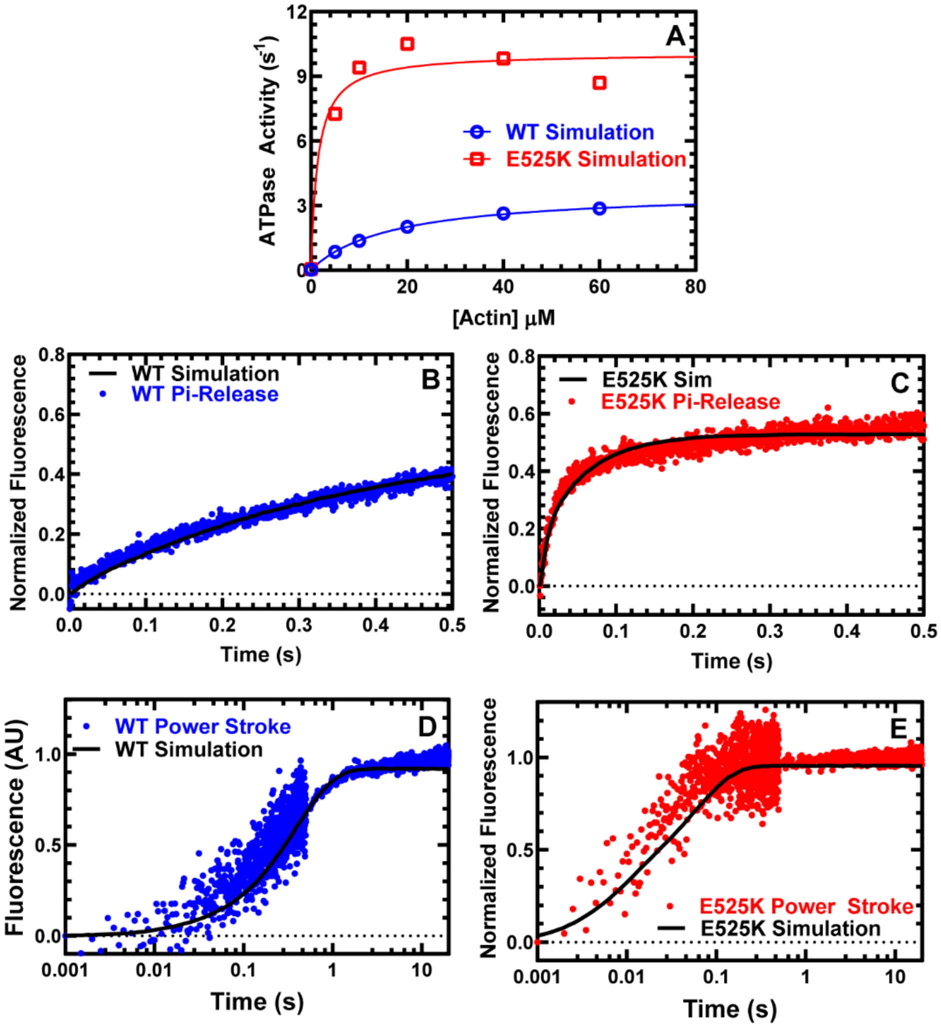


**Figure S5. Kinetic simulations. (A)** The rate and equilibrium constants determined experimentally and estimated from previous studies (**see Table S1**) were used to simulate the steady-state ATPase for both WT and E525K. **(B & C)** The phosphate release transients and **(D & E)** power stroke transients were compared to simulated data in WT and E525K, respectively. Only the first 0.5 sec of the Pi-release transients were simulated to focus on the initial Pi-burst associated with actin-activated Pi-release.

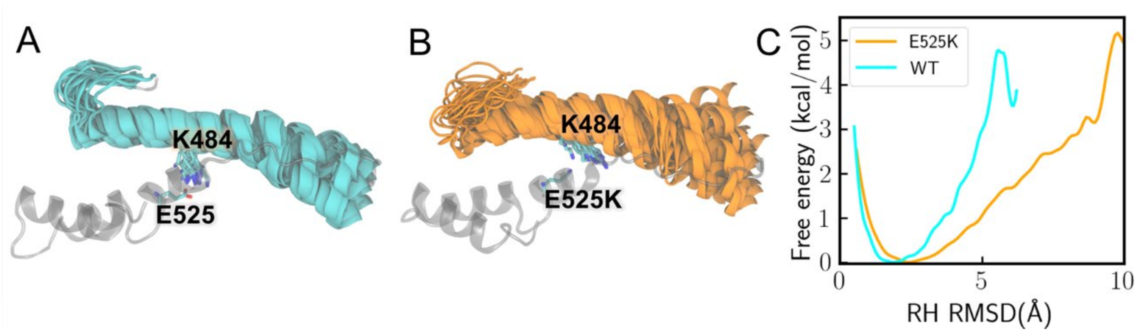


**Figure S6. Enhanced flexibility of the relay helix in the E525K simulations. (A) (B)** All the MD clusters of the relay helix (RH) are overlaid after being aligned to residues 519 to 540 (WT in cyan; E525K in orange). Variations in the K484 side chain are also demonstrated in both panels. **(C)** The free energy profile is plotted along the RH RMSD coordinate. E525K is much more likely than WT to exhibit RH RMSD values greater than 3Å.

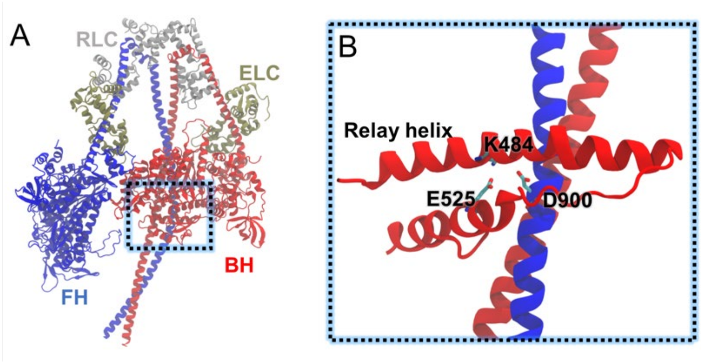


**Figure S7. Model to explain how E525K may stabilize IHM formation.** By forming an additional salt bridge (E525K and D900), the mutation E525K could stabilize the interface between the blocked head (BH-red) and subfragment-2 of the free head (FH-blue). The 8ACT PDB structure was used to generate the IHM model.

**Supporting Information References**


1. H. D. White, B. Belknap, M. R. Webb, Kinetics of Nucleoside Triphosphate Cleavage and Phosphate Release Steps by Associated Rabbit Skeletal Actomyosin, Measured Using a Novel Fluorescent Probe for Phosphate. *Biochemistry* **36**, 11828-11836 (1997).
2. A. M. Swenson *et al.*, Omecamtiv Mecarbil Enhances the Duty Ratio of Human beta-Cardiac Myosin Resulting in Increased Calcium Sensitivity and Slowed Force Development in Cardiac Muscle. *J Biol Chem* **292**, 3768-3778 (2017).
3. D. Chow, R. Srikakulam, Y. Chen, D. A. Winkelmann, Folding of the Striated Muscle Myosin Motor Domain. *Journal of Biological Chemistry* **277**, 36799-36807 (2002).
4. Q. Wang, C. L. Moncman, D. A. Winkelmann, Mutations in the motor domain modulate myosin activity and myofibril organization. *Journal of Cell Science* **116**, 4227-4238 (2003).
5. D. A. Winkelmann, E. Forgacs, M. T. Miller, A. M. Stock, Structural basis for drug-induced allosteric changes to human β-cardiac myosin motor activity. *Nature Communications* **6**, 7974 (2015).
6. W. Tang, J. Ge, W. C. Unrath, R. Desetty, C. M. Yengo, Cardiomyopathy mutations impact the actin-activated power stroke of human cardiac myosin. *Biophys J* **120**, 2222-2236 (2021).
7. D. V. Rasicci *et al.*, Impact of regulatory light chain mutation K104E on the ATPase and motor properties of cardiac myosin. *J Gen Physiol* **153** (2021).
8. J. D. Pardee, J. A. Spudich, Purification of muscle actin. *Methods Enzymol* **85 Pt B**, 164-181 (1982).
9. E. M. De La Cruz, E. Michael Ostap, "Chapter 6 Kinetic and Equilibrium Analysis of the Myosin ATPase". (Elsevier, 2009), 10.1016/s0076-6879(08)04206-7, pp. 157-192.
10. E. L. Smith, *Principles of Biochemistry, General Aspects* (McGraw-Hill, 1983).
11. D. V. Rasicci *et al.*, Dilated cardiomyopathy mutation E525K in human beta-cardiac myosin stabilizes the interacting-heads motif and super-relaxed state of myosin. *Elife* **11** (2022).
12. R. F. Sommese *et al.*, Molecular consequences of the R453C hypertrophic cardiomyopathy mutation on human β-cardiac myosin motor function. *Proceedings of the National Academy of Sciences* **110**, 12607-12612 (2013).
13. H. Miyata, R. Yasuda, K. Kinosita, Jr., Strength and lifetime of the bond between actin and skeletal muscle alpha-actinin studied with an optical trapping technique. *Biochim Biophys Acta* **1290**, 83-88 (1996).
14. T. ENDO, T. MASAKI, Molecular Properties and Functions In Vitro of Chicken Smooth-Muscle α-Actinin in Comparison with Those of Striated-Muscle α-Actinins1. *The Journal of Biochemistry* **92**, 1457-1468 (1982).
15. A. Suzuki *et al.*, Some properties of purified skeletal muscle alpha-actinin. *Journal of Biological Chemistry* **251**, 6860-6870 (1976).
16. M. J. Greenberg, J. R. Moore, The molecular basis of frictional loads in the in vitro motility assay with applications to the study of the loaded mechanochemistry of molecular motors. *Cytoskeleton* **67**, 273-285 (2010).
17. W. Ma, S. You, M. Regnier, J. A. McCammon, Integrating comparative modeling and accelerated simulations reveals conformational and energetic basis of actomyosin force generation. *Proceedings of the National Academy of Sciences* **120** (2023).
18. R. Salomon-Ferrer, D. A. Case, R. C. Walker, An overview of the Amber biomolecular simulation package. *WIREs Computational Molecular Science* **3**, 198-210 (2013).
19. J. A. Maier *et al.*, ff14SB: Improving the Accuracy of Protein Side Chain and Backbone Parameters from ff99SB. *Journal of Chemical Theory and Computation* **11**, 3696-3713 (2015).
20. Y. Miao, V. A. Feher, J. A. McCammon, Gaussian Accelerated Molecular Dynamics: Unconstrained Enhanced Sampling and Free Energy Calculation. *Journal of Chemical Theory and Computation* **11**, 3584-3595 (2015).
